# Transient glycan-shield reduction induces CD4-binding site broadly neutralizing antibodies in SHIV-infected macaques

**DOI:** 10.1101/2024.12.30.630768

**Authors:** Daniel J. Morris, Jason Gorman, Tongqing Zhou, Jinery Lora, Andrew J. Connell, Hui Li, Weimin Liu, Ryan S. Roark, Mary S. Campion, John W. Carey, Rumi Habib, Yingying Li, Christian L. Martella, Younghoon Park, Ajay Singh, Kirsten J. Sowers, I-Ting Teng, Shuyi Wang, Neha Chohan, Wenge Ding, Craig Lauer, Emily Lewis, Rosemarie D. Mason, Juliette M. Rando, Lowrey Peyton, Chaim A. Schramm, Kshitij Wagh, Bette Korber, Michael S. Seaman, Daniel C. Douek, Barton F. Haynes, Daniel W. Kulp, Mario Roederer, Beatrice H. Hahn, Peter D. Kwong, George M. Shaw

**Affiliations:** Departments of Medicine and Microbiology, Perelman School of Medicine, University of Pennsylvania, Philadelphia, PA 19104, USA; Vaccine Research Center, National Institute of Allergy and Infectious Diseases, National Institutes of Health, Bethesda, MD 20892, USA; Division of Viral Products, Center for Biologics Evaluation and Research, Food and Drug Administration, Silver Spring, MD 20993, USA; Aaron Diamond AIDS Research Center, Columbia University Vagelos College of Physicians and Surgeons, New York, NY 10032, USA; Vaccine and Immunotherapy Center, The Wistar Institute, Philadelphia, PA 19104, USA; Duke Human Vaccine Institute, Duke University School of Medicine, Durham, NC 27710, USA; New Mexico Consortium, Los Alamos, NM 87545, USA; Center for Virology and Vaccine Research, Beth Israel Deaconess Medical Center, Boston, MA 02215, USA; Department of Immunology, Duke University School of Medicine, Durham, NC 27710, USA; Duke Human Vaccine Institute, Duke University School of Medicine, Durham, NC 27710, USA; Department of Medicine, Duke University School of Medicine, Durham, NC 27710, USA

**Keywords:** broadly neutralizing antibody, CD4-binding site, co-evolution, glycan deletion, HIV-1, SHIV, structure-guided vaccine design

## Abstract

Broadly neutralizing antibodies (bNAbs) targeting the HIV-1 CD4-binding site (CD4bs) occur infrequently in macaques and humans and have not been reproducibly elicited in any outbred animal model. To address this challenge, we first isolated RHA10, an infection-induced rhesus bNAb with 51% breadth. The cryo-EM structure of RHA10 with HIV-1 envelope (Env) resembled prototypic human CD4bs bNAbs with CDR-H3-dominated binding. Env-antibody co-evolution revealed transient elimination of two Env CD4bs-proximal glycans near the time of RHA10-lineage initiation, and these glycan-deficient Envs bound preferentially to early RHA10 intermediates, suggesting glycan deletions in infecting SHIVs could consistently induce CD4bs bNAbs. To test this, we constructed SHIV.CH505.D3 with CD4bs-proximal glycan deletions. Infection of 10 macaques resulted in accelerated CD4bs bNAb responses in 8, compared with 1 of 115 control macaques. Glycan hole-based immunofocusing coupled to Env-Ab co-evolution can consistently induce broad CD4bs responses in macaques and thus serve as a model for HIV vaccine design.

**Highlights:**

- Out of 115 wildtype HIV-1 Env bearing SHIV infected macaques, only one macaque (T681) developed CD4bs bNAbs
- CD4bs bNAbs in macaque T681 recognized Env similarly to previously described CDR-H3 dominated human CD4bs bNAbs and exhibited comparable breadth and potency
- Transient CD4bs-proximal glycan deletions in macaque T681 preceded bNAb induction
- A novel SHIV with CD4bs-proximal glycan holes and enhanced CD4bs antigenicity immunofocused B cell responses to the CD4bs and elicited cross-clade neutralizing responses in 80% of macaques

## INTRODUCTION

A major goal of HIV-1 vaccine design is the consistent elicitation of broadly neutralizing antibodies (bNAbs). Reverse vaccinology approaches have sought to elucidate viral envelope-antibody (Env-Ab) co-evolutionary pathways that occur in HIV-1 infected humans who develop bNAbs to identify candidate priming and boosting immunogens that might replicate this co-evolution^1^. Studying the fine details of human bNAb development is challenging due to a paucity of longitudinally sampled bNAb subjects with cryopreserved blood mononuclear cells and plasma for analysis. Even in participants that are well sampled, only a minority are identified during early infection when transmitted/founder (T/F) envelopes can be inferred and Env-Ab coevolution fully evaluated^2^.

To address these limitations, we developed simian-human immunodeficiency viruses (SHIVs) bearing primary or T/F HIV-1 Envs that replicate persistently in rhesus macaques (RMs)^3^. These SHIVs contain a single mutation at Env residue 375 that promotes rhesus-CD4 binding and infection of rhesus CD4+ T cells while retaining antigenic features of the corresponding wild-type HIV-1 Env^3^. With this enabling technology, critical steps in Env-Ab coevolution can be analyzed from the time of virus infection to the development of autologous NAbs, and in some cases, heterologous bNAbs. Previously, we reported that macaques infected with SHIVs expressing HIV-1 Envs CH505, CH848, and CAP256SU recapitulated key features of viral evolution observed in humans from which these envelopes originated^4^. In addition, we found that in a subset of these SHIV-infected macaques, bNAbs developed that targeted the same canonical bNAb epitope supersites as in the corresponding human subjects, including the V2 apex and the V3 glycan patch^4^. Interestingly, in SHIV.CH505 infected macaques, we found evidence of VH-gene restricted, heavy chain complementarity determining region 2 (CDR-H2) mediated, CD4-binding site (CD4bs) targeted antibodies but these lacked neutralization breadth because of their exceptionally long light chain CDR-L1 of 17 amino acids^4^. Altogether, the rhesus antibodies shared phenotypic, genotypic, and structural similarities with the human mAbs that typify these bNAb classes.

Developing a vaccine to elicit bNAbs targeting the CD4bs epitope of Env has been a high priority due to the superior breadth and potency of these antibodies. CD4bs bNAbs identified from HIV-infected donors fall into two major classes: CDR-H3 dominated and VH-gene restricted (CDR-H2 mediated)^5^. While CDR-H3 dominated bNAbs can have breadths reaching 82%^5^, breadth from VH-gene restricted antibodies such as antibody N6 can have breadths reaching 98%^6^. Despite these promising features, vaccine-elicited CD4bs bNAbs have yet to be reproducibly induced in a standard outbred vaccine-test species. For CDR-H3 dominated CD4bs bNAbs^5,7^, Sanders and colleagues identified such antibodies, albeit with low breadth and potency, in BG505 GT1.1 SOSIP Env trimer vaccinated RMs^8,9^. For VH-gene restricted CD4bs bNAbs, substantial progress has been made in priming VRC01- or CH235-class antibodies^10–12^, which utilize heavy chain variable gene IGHV1-2*02 or IGHV1-46*01, respectively, and whose CDR-H2 mimics human CD4 in making contact with the CD4bs on Env. Furthermore, Saunders and colleagues recently identified rhesus CD4bs targeted antibodies that utilize the IGHV1-105 gene (KIMDB^13^), which is orthologous to the human IGHV1-46 gene (IGMT^14^) and mediates HIV-1 CD4bs binding by a CDR-H2 restricted mechanism^15^, but none of these antibodies exhibited neutralization breadth. Given the extensive affinity maturation required for breadth development^16^, it remains unclear how VH-gene restricted or CDR-H3 dominated CD4bs bNAbs might be consistently induced in humans or macaques.

In this context, we sought to determine if CD4bs bNAbs could be induced in SHIV-infected RMs as proof-of-feasibility for their induction by vaccination and potentially as a molecular guide for sequential immunogen designs. Previously, immunization with soluble Env trimers containing targeted deletions of four glycans proximal to the CD4-binding site consistently elicited CD4bs antibodies^17^; such antibody responses, however, could only neutralize viral strains that contained the same glycan deletions, and it was unclear how to mature these responses to enable neutralization of fully glycosylated heterologous viral strains. To gain insight into how CD4bs antibodies capable of neutralizing wild-type heterologous virus strains might be induced, we investigated the development of a potent, broadly neutralizing rhesus CD4bs antibody that resulted from persistent infection by a SHIV bearing a primary, wild-type T/F Env. Importantly, in this investigation, we determined that a CDR-H3 dominated bNAb with structural features and paratope-epitope interactions similar to human CD4bs bNAbs was induced. Surprisingly, in that macaque, we observed temporal linkage between spontaneously occurring glycan deletions surrounding the CD4bs in the evolving viral quasispecies and the initiation of the CD4bs bNAb lineage. We thus hypothesized that such glycan deletions, if incorporated prospectively into an infecting SHIV, might enhance the critical priming step in bNAb induction and then serve to immunofocus and affinity-mature bNAb development by persistent virus replication and Env-Ab co-evolution. In this study, we tested this hypothesis by using structure-based designs to create glycan-modified SHIVs and assessed their ability to induce CD4bs bNAb responses in RMs.

## RESULTS

### SHIV.CH1012-infected macaque T681 developed a broadly neutralizing antibody lineage RHA10 targeting the CD4-binding site

We constructed SHIV.CH1012 from the T/F Env inferred from an acute infection cohort study participant, CH1012, as previously described^18^ and infected six Indian-origin rhesus macaques (**Figure S1A**). Animals were bled longitudinally to quantify plasma vRNA and monitor neutralization activity (**Figures S1B, S1C**). All macaques exhibited typical acute replication kinetics, with plasma viral RNA titers of ∼10^7^ mRNA/ml two weeks post-SHIV inoculation (wpi). All six macaques developed neutralizing antibodies against the autologous tier-2 SHIV.CH1012 virus and against several tier-1a viruses (**Figure S1C**). We screened for heterologous tier-2 neutralization in these animals and found that only RM T681 developed broadly neutralizing plasma antibodies beginning at 32 wpi, which increased in breadth and potency over time reaching 55% breadth against a screening panel of 18 tier 1B/2 heterologous viruses (**Figures 1A, S1C**). T681 plasma exhibited differential binding to RSC3^19^ and BG505 DS-SOSIP^20^ probes with CD4bs KO mutants suggesting the presence of binding antibodies to the CD4bs at a timepoint corresponding to broad neutralization (**Figure 1B**). We next tested plasmas against a panel of BG505.T332N mutant pseudoviruses to map the epitope of this broad neutralization. T681 plasma neutralization of BG505 at multiple time points was enhanced against glycan 363 or glycan 462 knockout mutants (**Figure 1C**) but not against glycan 276 knockout mutants. This phenotype suggested CD4bs targeting, but in a manner different from human CD4bs bNAbs VRC01, CH235, and CH103 that are primarily enhanced by N276 glycan removal (**Figure 1C**).

**Figure 1.**
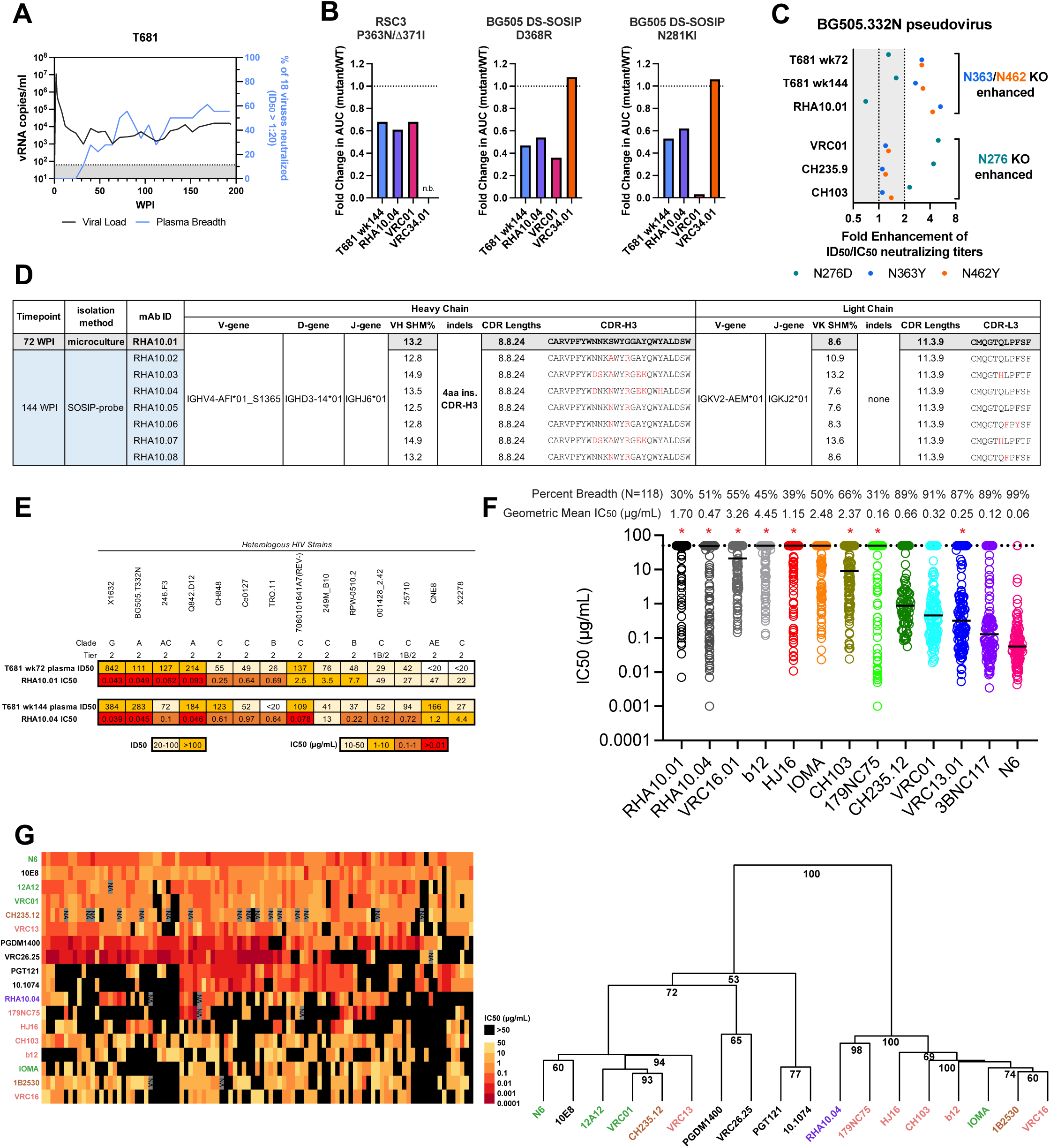
RHA10 is a rhesus CD4 binding site broadly neutralizing antibody with 51% neutralization breadth that is N276-glycan agnostic. (A) Longitudinal plasma vRNA load (lower limit of quantification: 62 RNA copies/mL) and percent heterologous breadth of an 18-strain panel in macaque T681 (B) Fold change in binding AUC between WT and CD4bs knockout mutant RSC3 cores or BG505 DS-SOSIPs, n.b. – no binding to WT protein (C) Fold change in neutralization titers of T681 plasma or mAbs against BG505.332N pseudovirus glycan knockout mutants. Enhancement is defined as a greater than 2-fold increase in ID50/IC50 titers. (D) Immunogenetics summary of RHA10 lineage, RHA10.01 highlighted as a reference with further SHM in V(D)J junction in red (E) Recapitulation of T681 plasma neutralization breadth and potency by respective RHA10 lineage members. (F) Neutralization breadth and potency on a 118-virus panel of RHA10 members and human CD4bs bNAbs tested on the panel. Potency and breadth values are listed above each mAb. Median IC50 titers are represented as black bars. Human CD4bs bNAb neutralization data obtained from CATNAP. (G) Heatmap and hierarchical clustering of large pseudovirus panel neutralization data. Only bootstrap values above 50% are shown. CD4bs bNAbs colored by class, RHA10.04 (purple), CDR-H3 dominated (salmon), VH1-2 restricted (green), and VH1-46 restricted (brown).

To confirm this CD4bs bNAb specificity and elucidate the molecular basis of the broad neutralization in T681, we isolated broadly neutralizing mAbs by sorting IgG+ memory B cells from peripheral blood mononuclear cells (PBMCs) at 72 and 144 wpi using an unbiased B cell culture and SOSIP-sorting strategy, respectively^21^ (**Figure S1D**). From these sorts, we identified RHA10, a lineage of antibodies utilizing *Macmul* IGHV4-AFI*01_S1365 with a 24 amino acid (a.a.) long CDR-H3 and IGKV2-AEM*01 with a 9 a.a. CDR-L3 (**Figures 1D, S1D**). These mAbs recapitulated the breadth and potency of the plasma antibodies from the corresponding time points (**Figures 1E, S1C**). RHA10.04 exhibited a similar binding profile to T681 wk144 plasma (**Figure 1B**) and RHA10 members exhibited heterologous neutralization of BG505, which was enhanced by glycan 363 and 462 knockouts but were unaffected by glycan 276 knockout (**Figure 1C**). Together these data suggested that RHA10 lineage antibodies were the sole contributor of breadth in macaque T681. The broadest mAb lineage member from each timepoint (RHA10.01 and RHA10.04) was tested against a 119-strain virus panel to better assess breadth and potency (**Figure 1F, Table S1**). RHA10.04 neutralized 51% of heterologous viruses with a 50 µg/mL cutoff at a geometric mean IC50 of 0.436 µg/mL. These values were comparable to other human CD4bs bNAbs that have been tested on the same panel^22^, particularly those that are CDR-H3 dominated and not restricted by variable heavy chain alleles VH1-2 or VH1-46. Hierarchical clustering of heterologous neutralizing IC50 titers showed that RHA10.04 robustly clustered with CD4bs bNAbs that had moderate breadth, with the closest being the CDR-H3 dominant binder 179NC75 (**Figure 1G**). In contrast V2 apex, V3 glycan, and MPER bNAbs clustered separately, as did the set of CD4bs bNAbs with the greatest breadth. Finally, RHA10.04 showed significantly correlated IC50 titers and sensitive/resistant patterns against large panels of pseudoviruses compared with prototypical CDR-H3 dominant CD4bs bNAbs including HJ16 and 179NC75 (**Figure S2)**. Thus, RHA10 is the first example, to our knowledge, of a SHIV- or vaccine-induced rhesus CD4bs antibody that is comparable to prototypic human CD4bs bNAbs and that reaches neutralization breadth and potency thresholds expected for effective clinical protection against HIV-1 challenge^23–25^.

### Structure of RHA10 with HIV-1 Env trimer revealed similarities to human CD4bs bNAbs HJ16 and 179NC75

We next used single particle cryogenic-electron microscopy (cryo-EM) to determine the structure of rhesus antibody RHA10.01 bound to the soluble pre-fusion stabilized envelope trimer BG505 DS-SOSIP at a resolution of 4.3Å (**Figures 2A, S3, Table S2**). RHA10.01 bound the trimer at a 3:1 stoichiometry at the CD4bs, but at an angle distinct from other human CD4bs bNAbs. Despite its unique angle of approach, the envelope footprint of RHA10.01 overlapped with VRC01 and human CD4 (**Figures 2B, 2C**). The CDR-H3 of RHA10.01 contacted primarily conserved residues within the CD4bs (283, 457, 470, 474) likely contributing to its breadth (**Figure 2D**). The tip of RHA10.01’s CDR-H3 curled back toward the antibody core and made additional contacts with conserved residues in the CD4 binding loop (365, 366) and beta-sheet β24 (470) (**Figure 2E**) The angle of approach of RHA10.01 was also consistent with the glycan knockout phenotypes we observed previously (**Figure 1C**). RHA10 engaged the CD4bs from a similar latitudinal angle of approach as other CDR-H3 dominated human CD4bs bNAbs (HJ16, VRC13, VRC16, CH103) and CD4^5^ **(Figure 2F**). RHA10, however, approached the epitope from a longitudinal angle that was further away from the trimer axis (**Figure 2F**). Together this approach angle explains why RHA10 remains unaffected by changes to the N276 glycan (**Figure 1C, S3G**).

**Figure 2.**
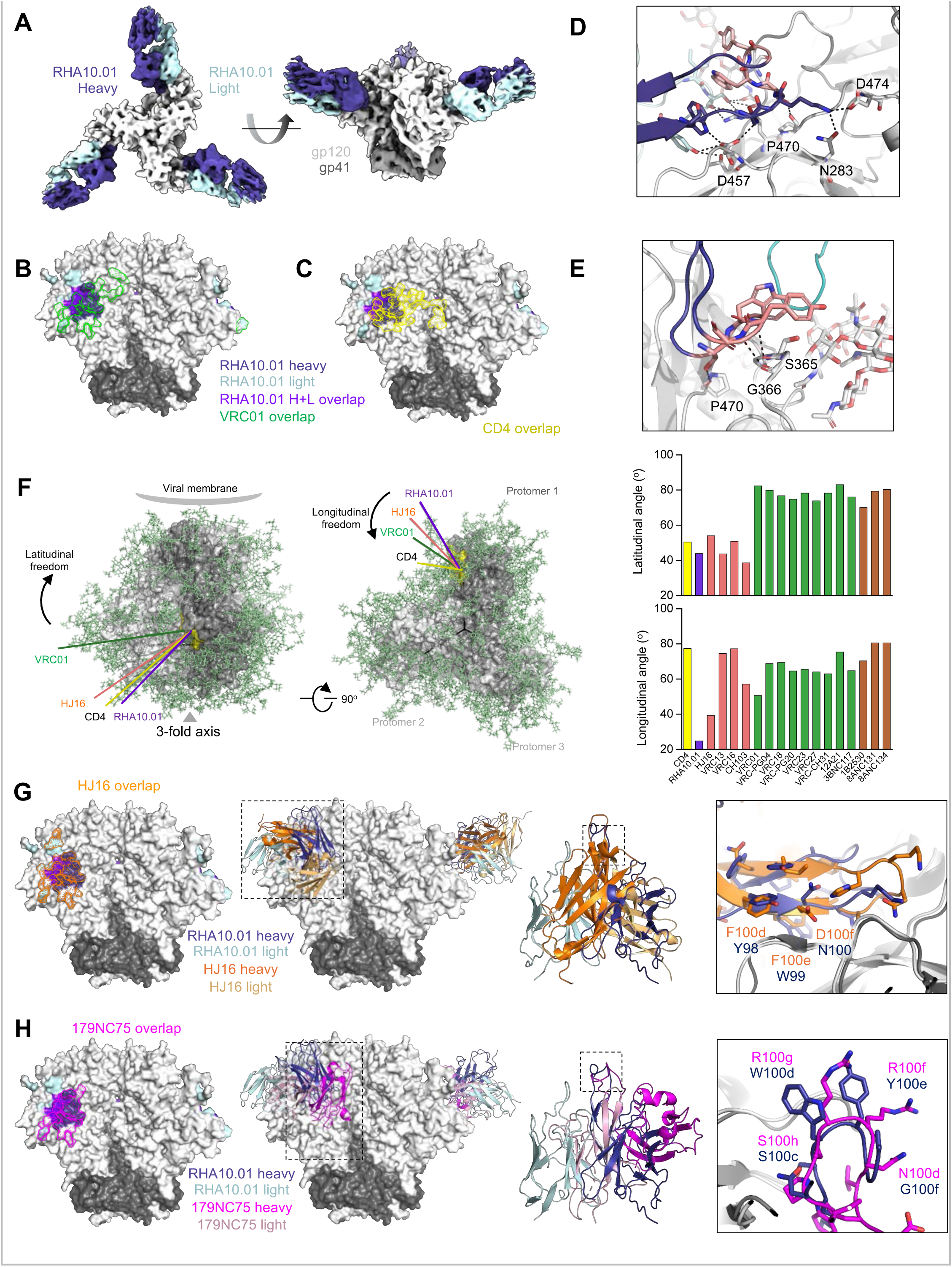
RHA10 binds the CD4-binding site primarily through its long CDR-H3. (A) Cryo-EM structure of RHA10.01 bound to BG505-DS-SOSIP, colored by chain (B) RHA10.01 binding footprint overlaid by VRC01 binding footprint (green) (C) RHA10.01 binding footprint overlaid by CD4 footprint (yellow) (D) RHA10.01 CDR-H3 stem Env contact details. CDR-H3 insertion colored in pink. (E) RHA10.01 CDR-H3 tip Env contact details. CDR-H3 insertion colored in pink. (F) Angle of approach analysis, CDR-H3 dominated class (salmon) VH1-2 class (green) VH1-46 class (brown) (G) RHA10.01 and HJ16 structural comparison by footprint, ribbon diagram overlap, and CDR-H3 geometries (H) RHA10.01 and 179NC75 structural comparison by footprint, ribbon diagram overlap, and CDR-H3 geometries

The heterologous virus neutralization profile of RHA10 was statistically similar to CDR-H3 dominated binders HJ16^26^ and 179NC75^27^ (**Figures 1G, S2**). From an alignment of the gp120 cores from both structures (HJ16 PDB ID: 4EY4, 179NC75 PDB ID: 7LLK), we see that the epitope footprints of RHA10.01 and HJ16 overlap well, although the heavy and light chain orientations between the two mAb differ substantially (**Figure 2G**). Although these orientations are reversed, we found an interesting overlap in the CDR-H3 stems of both mAbs. This alignment is particularly good for a _98_YWN_100_ motif in RHA10 and a _100d_FFD_100f_ motif in HJ16. These residues interact with CD4bs residues 457-459 which are highly conserved across strains. 179NC75 and RHA10.01 footprints also shared a large degree of overlap (**Figure 2H**). 179NC75 had a similar heavy-light chain orientation to RHA10 but overall little alignment. Interestingly, the CDR-H3 tips of both mAbs had a similar curl. Despite this backbone similarity, the CDR-H3 tip side chains varied substantially in composition and orientation between these two mAbs. These microhomologies likely explain why RHA10, HJ16, and 179NC75 were similar in the neutralization signature analysis although their angles of approach and the heavy-light chain orientations varied. These differences between RHA10, HJ16, and 179NC75 likely contributed to their different sensitivities to a glycan at residue 276^26–28^. Neutralization signature analysis and structure together demonstrated that RHA10 is similar to but distinct from all known human CD4bs bNAbs.

### RHA10 lineage acquired a four amino acid insertion by somatic hypermutation that was critical to breadth development

To define critical steps in RHA10 development from the initial naïve B cell priming event through affinity maturation to breadth and potency, we performed next-generation sequencing of bulk IgG+ memory B cells from longitudinal samples of PBMCs from RM T681. We also used IgDiscover^29^ to generate an animal-specific germline Ig-expressed gene allele dataset from sorted naïve B cells. Then, using SONAR^30^ and the two sequence datasets, we phylogenetically identified the RHA10 lineage and inferred its unmutated common ancestor (UCA) with high confidence (**Figures 3A, S4A, S4B, Table S3**). The RHA10 lineage was derived from *Macmul* IGHV4-AFI*01_S1365, IGDH3-14*01, IGHJ6*01 and *Macmul* IGKV2-AEM*01, IGKJ2*01 alleles for the heavy and kappa chains, respectively. The RHA10 UCA inference for heavy and light chains was unambiguous, revealing a 20 a.a. CDR-H3 that subsequently acquired a 4 a.a. insertion between 18-24 wpi (**Figures 3A, S4A**). B cells containing this 4 a.a. insertion were positively selected such that all RHA10 members identified by 40 wpi and later contained this insertion (**Figures 3B, S4A**). Finer inspection of early members of different RHA10 sublineages revealed alternative insertion events in different reading frames of the CDR-H3 in several sublineages of the RHA10 tree but these failed to gain full penetrance in later RHA10 lineage sequences (**Figure S4C**).

**Figure 3.**
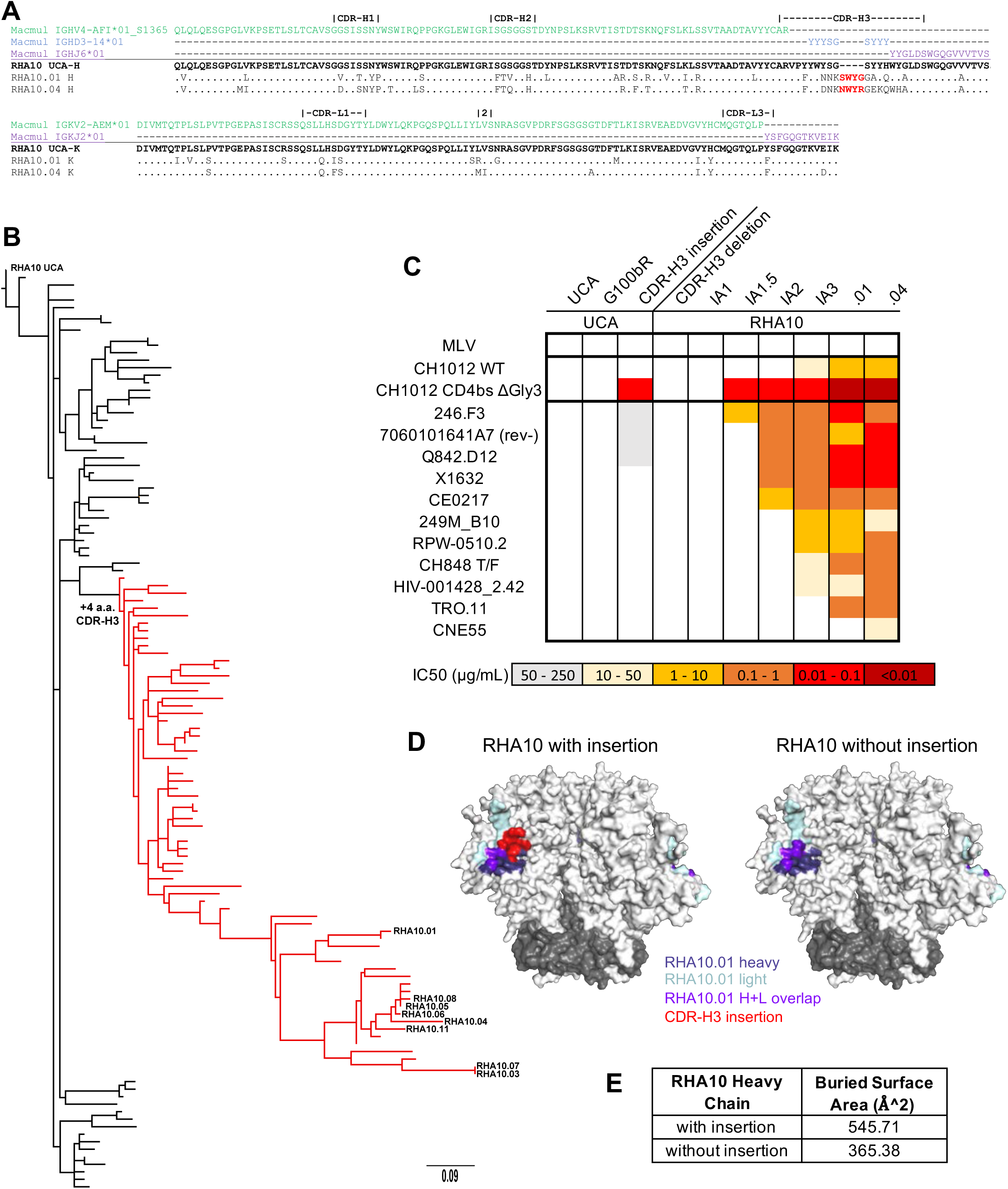
RHA10 developed a CDR-H3 insertion during maturation crucial for broad neutralization. (A) Amino acid alignments of RHA10.01 and RHA10.04 heavy and light chain genes to inferred germline genes. Dots represent identity to the unmutated common ancestor (UCA). “2”, CDR-L2 of three amino acids. (B) Phylogenetic tree of RHA10 lineage heavy chains rooted to the RHA10 UCA. Members containing the CDR-H3 insertion are highlighted in red. (C) Neutralization titers of RHA10 lineage evolutionary members and recombinant mutants against autologous and heterologous viral strains. CH1012 CD4bs ΔGly3 lacks the N197, N362, and N461 glycans. (D) Binding footprint comparison of RHA10.01 with and modeled without the CDR-H3 insertion (E) Buried surface area of RHA10 heavy chains as calculated by PISA.

We evaluated the importance of the 4 a.a. insertion that gave rise to the mature bNAb lineage members in two ways. First, we deleted the insertion from the mature bNAb RHA10.01 and assessed the impact on neutralization breadth and potency (**Figures 3C, S4D**). RHA10.01 neutralized numerous heterologous viruses as well as the autologous virus CH1012, whereas the RHA10 UCA neutralized none of these viruses even at concentrations as high as 250 μg/mL. Deleting the 4 a.a. insertion from the mature RHA10.01 antibody reverted its neutralization to a UCA-like phenotype. Second, we inserted the 4 a.a. sequence into the CDR-H3 of the RHA10 UCA, which expanded the breadth and potency of neutralization (**Figures 3C, S4D**). We also tested the effect of a G100_b_R heavy chain mutation as this was acquired before the insertion event. This substitution did not enhance RHA10 UCA neutralization of autologous or heterologous viruses. The CDR-H3 insertion into the RHA10 UCA was not sufficient to fully recapitulate RHA10.01 neutralization, indicating that additional somatic hypermutations contributed to the full breadth of the lineage. This was evident by testing the phylogenetically inferred RHA10 intermediate antibodies (RHA10 IA1-IA3) (**Figures 3C, S4D**). RHA10 IA3 contains the CDR-H3 insertion as well as additional SHM and almost completely recapitulated RHA10.01 neutralization whereas RHA10 IA1.5 and RHA10 IA2, which also contain the insertion but fewer SHMs, did not (**Figures 3C, S4D**). In the cryo-EM structure, the CDR-H3 insertion makes important contacts with Loop D, the CD4-binding loop, and ß24 (**Figures 2D, 2E**). By modeling the buried surface area of the RHA10.01 heavy chain with and without the 4 a.a. insertion, we could show that the insertion contributed nearly one-third of the heavy chain’s interaction with Env (**Figures 3D, 3E**). Overall, our results highlight the critical importance of a somatically acquired indel to the breadth and potency of RHA10, consistent with observations that other human CD4bs bNAb lineages also develop indels that contribute importantly to neutralization breadth and potency^31^.

### Transient deletion of glycans around the CD4-binding site coincided with RHA10-lineage initiation

Having established the potential for macaques to develop CD4bs bNAbs, we sought to explore how the evolving viral quasispecies in T681 might have contributed to RHA10 induction and affinity maturation. We first performed longitudinal single genome sequencing (SGS)^2^ of full-length *env* gp160 genes from plasma virus beginning at 4 wpi and extending to 160 wpi (**Figure S5A**). SGS proportionately represents target RNA molecules in a sample and allows genetic linkage to be maintained across full-length gp160 genes, thus facilitating analysis of Env phylogeny, function, antigenicity, and structure^2^. Phylogenetic analysis of gp160 sequences in plasma revealed early random nucleotide substitutions typical of acute HIV-1 infection^2^, followed by strong selection for nonsynonymous substitutions resulting in 2.86%-per year a.a. divergence and maximum diversity of 9.74% by three years. These substitutions were annotated in a *Pixel* plot (**Figure S5B**) which revealed positive selection at potential N-linked glycosylation sites (PNGS) and at predicted NAb and cytotoxic T-cell epitopes across Env. Surprisingly, we observed that near the time of RHA10 elicitation at 18 wpi, 20% of the sequences had lost the PNGS at residue 362, and 50% had lost the PNGS at residue 461 (**Figure 4A**). The loss of these glycans was transient, as sequences at 24/32 wpi with the PNGS at 362 and 461 restored were detected, coincident with the first detection of plasma breadth. These changes were of particular interest to us since they correlated temporally with the emergence of RHA10 bNAb lineage in the memory B cell population (**Figure S4A**) and deletion of the same PNGS in Env trimer immunogens had, in earlier studies, strongly immunofocused antibody responses to the CD4bs^10,17,32–34^. To more precisely estimate the prevalence and temporal occurrence of PNGS deletions at residues 362 and 461 in the evolving T681 viral quasispecies, we performed deep next-generation sequencing (NGS) of an amplicon spanning both PNGS in plasma viral RNA from serial time points. Sequencing depth of these NGS analyses was 74,848-161,510 sequences compared to <100 SGS sequences. The NGS results confirmed that PNGS at positions 362 and 461 were disrupted just before RHA10 elicitation, with 50% of the evolving viral quasispecies at 18 wpi lacking a PNGS at either position and approximately 5% of NGS reads lacking PNGS at both positions (**Figure 4A, Table S4**). This temporal concordance suggested the possibility that circulating viruses lacking one or both of these glycans may have bound the RHA10 UCA and triggered the lineage.

**Figure 4.**
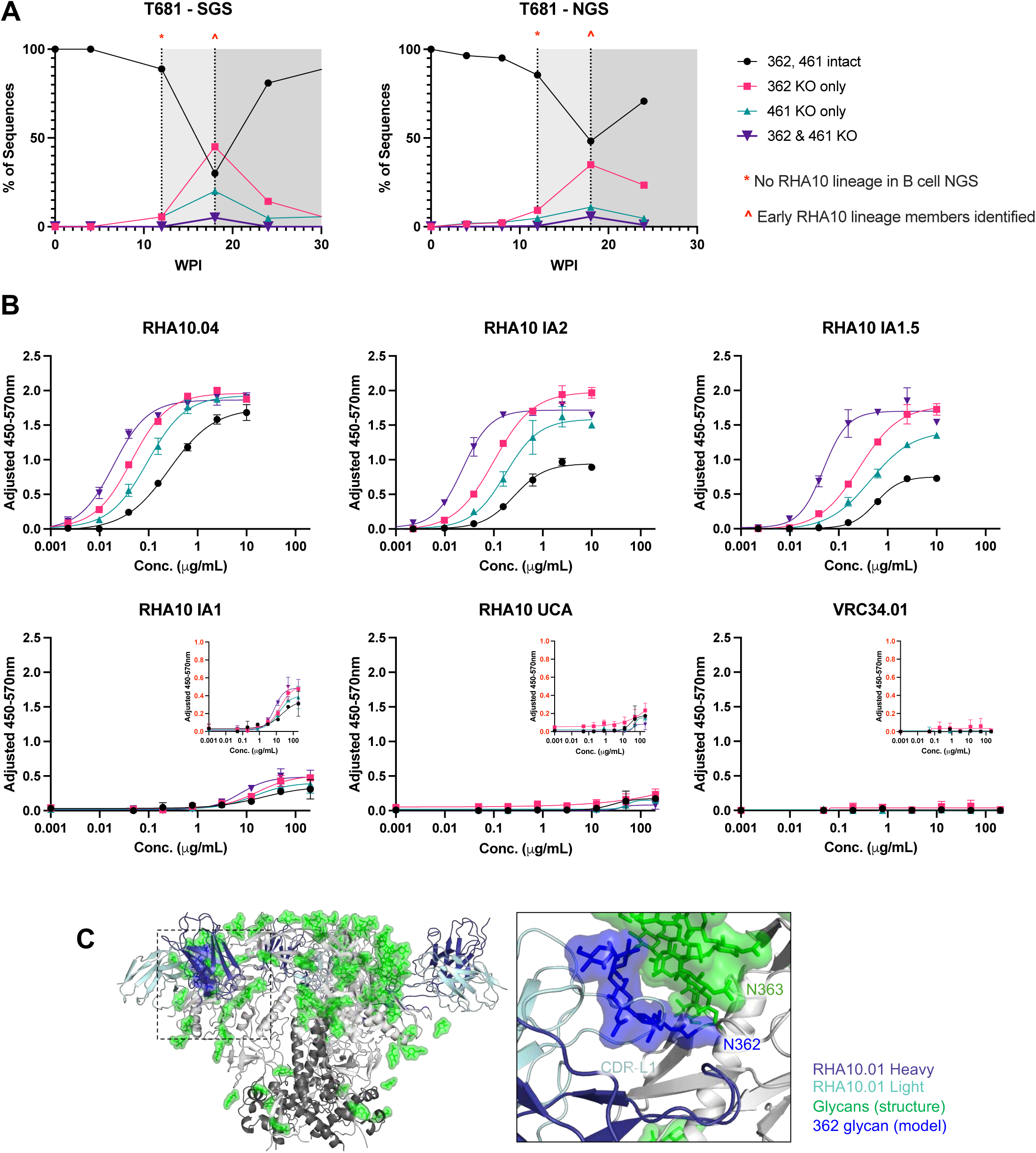
Transient deletion of glycans surrounding the CD4bs preceded RHA10 elicitation. (A) Percentage of single genome viral sequences (left) or next-generation sequences (right) with mutated PNGS at residues 362 and/or 461. Dotted vertical lines represent important time points in RHA10 lineage development highlighted in the text. WPI, weeks post-infection. (B) ELISA curves of CH1012 gp120s binding by RHA10 UCA, inferred intermediates, RHA10.04, and negative control VRC34.01. Error bars represent standard deviation from the mean. (C) RHA10.01 structure from with glycans highlighted. N362 glycan present in CH1012 (blue) modeled in next to N363 glycan present in BG505 (green) clashing with RHA10.01 CDR-L1. 362 glycan model is based on gp120 alignment with PDB ID 5FYK.

To determine if RHA10 lineage antibodies bound preferentially to Envs devoid of glycans at positions 362 and 461, we tested their binding to recombinant CH1012 T/F Envs (gp120) lacking these glycan motifs (**Figures 4B, S5C**). By ELISA, we saw that the mature antibody RHA10.04 and its inferred ancestors IA2, IA1.5, and IA1 all showed enhanced binding to the glycan-deficient gp120s compared to wild-type Env (**Figure 4B**). Envs lacking both glycans exhibited increased binding compared to gp120s lacking just one glycan. RHA10 UCA did not show binding to any Envs, with or without the glycan deletions. We modeled the position of the 362 glycan in CH1012 Env against the 363 glycan present in the BG505 DS-SOSIP structure and found a predicted clash between the CH1012 N362 glycan and the RHA10.01 CDR-L1 (**Figure 4C**), which offered a structural explanation for preferred binding of glycan deleted CH1012 Envs by early RHA10 intermediates. By affinity maturation, the RHA10 lineage acquired a mutation at residue 32 in CDR-L1 where an aspartic acid to serine substitution is predicted to reduce CDR-L1 clashing with the N362 glycan (**Figure S4D**). To further assess RHA10 UCA binding to CH1012 glycan knockouts, we performed biolayer interferometry (BLI) with RHA10 UCA and RHA10.04 antibodies **(Figure S5D**). K_D_ values for the mature RHA10.04 roughly mirrored the ELISA data with an affinity to CH1012 T/F gp120 of 34.48 nM that was enhanced to 20.09 nM by an N362 glycan knockout, and 11-fold by the double glycan knockout gp120. RHA10 UCA binding to the gp120s was too low to accurately estimate binding affinities. Altogether, these data suggested that transient deletion of glycans surrounding the CD4bs may have triggered the RHA10 lineage in macaque T681.

### SHIVs lacking CD4bs-proximal glycans immunofocus antibody responses to the CD4bs in infected macaques

Previous studies have explored glycan deletion or glycan under-processing to immunofocus B cell responses to the underlying conserved protein surface as a strategy to prime CD4bs bNAb precursors^10,11,33–37^. However, these approaches have generally lacked a means to then affinity-mature on-target responses to acquire neutralization breadth other than to boost with Env trimers with restored glycans. We hypothesized that SHIVs containing structure-guided glycan deletions surrounding the CD4bs would prime B cell responses similarly, and then by progressively restoring the missing glycans and mutating additional sites in the epitope supersite, immunofocus and shepherd a subset of B cell lineages specifically to the glycan-shielded CD4bs. A key difference between this approach and previous strategies using Env trimer immunogens is that the SHIV “immunogen” replicates persistently and coevolves with B cell responses, thereby offering on-target bNAb lineage B cells a continuous affinity gradient. A second difference is that we chose to retain the N276 glycan in our SHIV designs to prime and then “train” B cell responses to accommodate this critical glycan that is present in most wild-type viruses and represents a developmental barrier for most CD4bs bNAbs^28,38^. Thus, we constructed SHIVs with glycan deletions at positions 362 and 461 (termed “D2”) or at positions 197, 362, and 461 (termed “D3”) in three different SHIV Env backbones (BG505.T332N, CH505 T/F and CH1012 T/F) (**Figure 5A**). Additionally, we designed SHIV.BG505.S241N.D3 that fills the 241 glycan hole that BG505 naturally contains. We and others have shown that the 241/289 glycan hole is an immunodominant, strain-specific antibody target during BG505 immunization or SHIV infection^39–43^.

**Figure 5.**
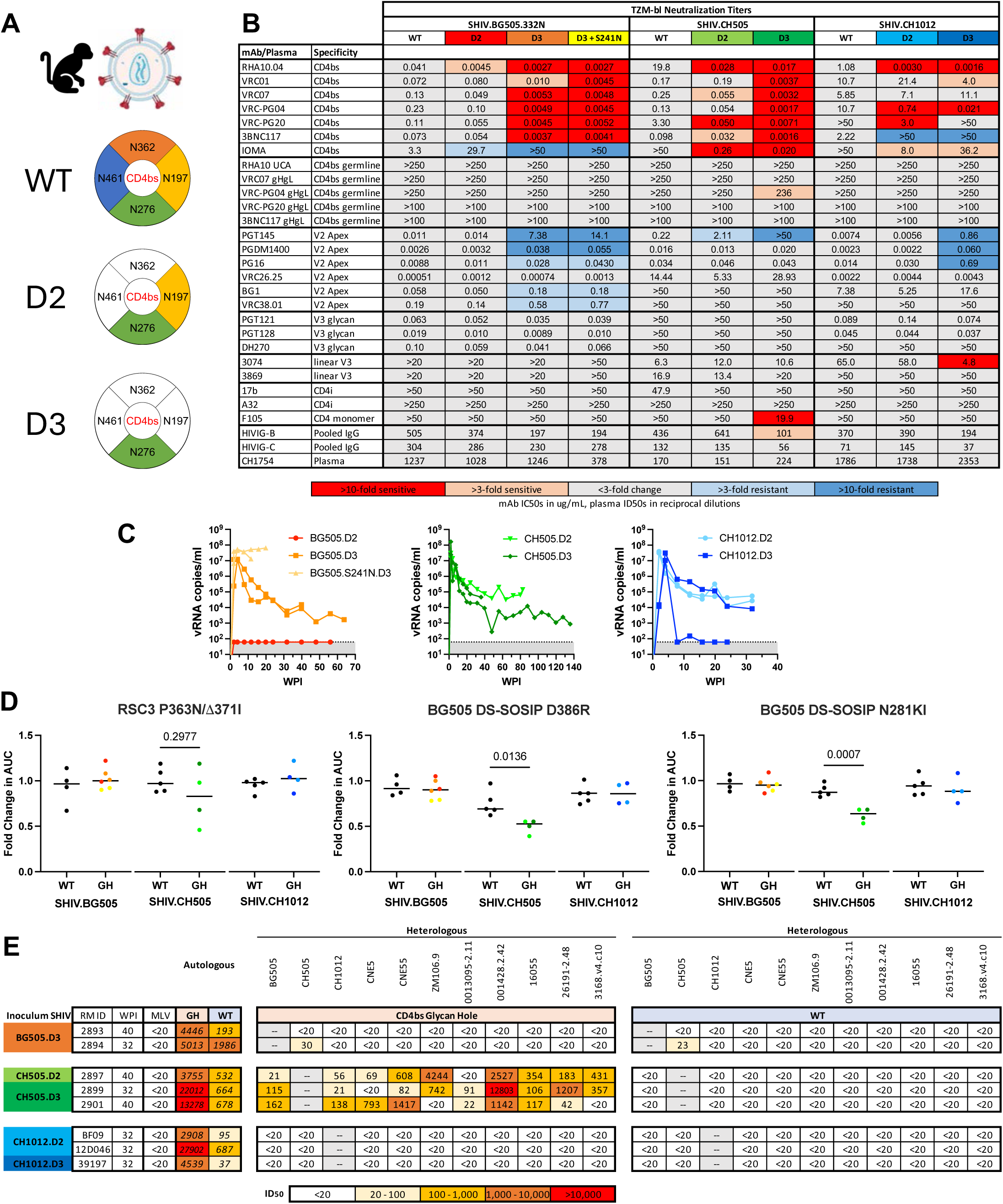
CD4bs glycan-deficient SHIVs improved CD4bs antigenicity, maintained infectivity, and elicited improved CD4bs targeting. (A) CD4bs glycan knockout design schematic, created in Biorender.com. D2 designs lack the N362 and N461 glycans. D3 designs lack the N197, N362, and N461 glycans. (B) Neutralization titers of CD4bs glycan-deficient SHIV designs, IC50 fold change highlighted (C) Viral load kinetics of CD4bs.GH SHIV-infected RMs, lower limit of quantification = 62 vRNA copies/mL. WPI, weeks post-infection. (D) Fold change in binding AUC of rhesus plasma between WT and mutant RSC3 and BG505 DS-SOSIP proteins. Unpaired t-tests were used for comparing WT and GH groups. (E) Plasma ID50s of autologous and heterologous viruses with or without the CD4bs glycans from CD4bs.GH SHIV-infected RMs.

We tested the antigenicity of these SHIV designs *in vitro* on TZM cells against a panel of neutralizing and non-neutralizing mAbs, inferred CD4bs bNAb precursors, and polyclonal anti-HIV-1 plasma (**Figure 5B**). As expected, we found that deleting these glycans increased the neutralization potency of CD4bs bNAbs including RHA10.04 as much as 1,000-fold. SHIV.CH505.D3 had increased sensitivity to all CD4bs bNAbs tested and was sensitive to the germline-reverted construct of human CD4bs bNAb VRC-PG04. Sensitivity to V3-glycan bNAbs and non-neutralizing mAbs was generally not affected more than 3-fold. Interestingly, V2 apex bNAbs, which are sensitive to the quaternary structure of the apex, did show reductions in their neutralization titers against D3 constructs, suggesting some disruption of the overall integrity of the Env trimer. However, because we did not observe large increases in CD4i, linear V3, or HIVIG neutralization titers against these constructs, we concluded that the D2 and D3 designs did not substantially alter the tier-2 neutralization phenotype of these Envs.

As a pilot study, we inoculated a small number of macaques to determine *in vivo* replication efficiency of these SHIVs as well as their persistence over time. Overall, we found that these SHIVs maintained wild-type levels of infectivity, replication efficiency and persistence, as inferred by plasma viral load profiles at early (2-4 wpi), intermediate (8-24 wpi), and late (>24 wpi) stages of infection (**Figure 5C**), although there were exceptions: macaques infected with SHIV.BG505.S241N.D3 exhibited exceptionally high peak and set point viral loads and rapidly progressed to AIDS, whereas one macaque infected with SHIV.CH1012.D3 controlled virus replication after 8 weeks^44^. All CD4bs.GH pilot RMs, as well as RMs infected with WT versions of SHIV.BG505, CH505, and CH1012 were screened for differential binding to RSC3, BG505 DS-SOSIP, and corresponding CD4bs KO proteins to identify on-target responses (**Figure 5D**). RMs infected with SHIV.CH505-based glycan hole designs exhibited statistically significant differential binding to wild-type and KO probes compared to sera from wild-type SHIV.CH505-infected macaques. RMs infected with SHIV.BG505 or SHIV.CH1012-based glycan hole designs did not show differential CD4bs binding in their plasma. We also screened RMs for autologous and heterologous neutralization against a virus panel containing wild-type and CD4bs glycan KO mutants. We found that all macaques developed autologous neutralization that was greatly enhanced by knocking out the CD4bs glycans, suggesting on-target CD4bs-focused responses (**Figure 5E, Table S5**). However, only macaques infected with SHIV.CH505-based designs developed antibody responses capable of neutralizing heterologous viruses lacking the CD4bs glycans, although these early sera could not neutralize wild-type viruses. These findings suggested that the early antibody responses elicited by these glycan hole engineered SHIVs recognized the conserved underlying epitopes of the CD4bs but that they could not accommodate wild-type glycans. Thus, we next asked: Could SHIV co-evolution, by filling glycan holes and immunofocusing the maturing B cell response, drive the development of broadly neutralizing responses?

### SHIV.CH505.CD4bs.GH consistently elicits broad HIV neutralization targeting the CD4bs

To examine the reproducibility of CD4bs immunofocusing and bNAb elicitation, we infected eight additional RMs with SHIV.CH505.D3. Viral replication in this repeat cohort was consistent, leading to plasma viral load setpoints in most animals of 10^3^-10^5^ vRNA copies/mL (**Figure 6A**). This resulted in similar CD4bs-targeting binding antibody responses as in the pilot RMs (**Figure 6B**). We next compared plasma samples from RMs infected with wild-type SHIV.CH505.T/F^3,4^ to our SHIV.CH505.D3-infected RMs for CD4bs targeted binding, neutralizing breadth, and potency (**Figures 6B, 6C, 6D**). RMs infected with SHIV.CH505.D3 showed significantly reduced binding to both BG505 DS-SOSIP CD4bs knockout mutants at magnitudes comparable to RHA10-containing T681 plasma and the control mAbs RHA10.04 and VRC01. RSC3 P363N/Δ371I did not reveal significantly reduced binding by plasma antibodies from the SHIV.CH505.D3-infected macaques, although a subset of four macaque plasmas did show reductions similar to VRC01 and RHA10.04. Importantly, SHIV.CH505.D3-infected macaques exhibited substantially greater neutralization breadth and potency against the CD4bs glycan KO viruses than did SHIV.CH505.WT-infected RMs (**Figure 6C, Table S6**). In addition, there was a trend for SHIV.CH505.CD4bs.GH-infected macaques to have greater wild-type breadth at 40 wpi compared to wildtype-infected macaques. We continued to monitor these RMs for improvements in wild-type heterologous breadth on a 34-virus panel through 112 wpi. We found that 9 of 11 RMs developed neutralization of at least 5 of 34 isolates (15%) compared with none of 8 SHIV.CH505.WT-infected RMs or with only 1 in 115 RMs infected by any one of 16 SHIVs expressing different primary HIV-1 Envs^3,4,18,45^ (R.H. and G.M.S. *submitted*) (Fisher’s exact test, p<0.001 and p<0.0001, respectively). A subset of SHIV.CH505.CD4bs.GH-infected macaques (CN37, 2901, 2897, CN17, CL84) neutralized between 20 of 34 isolates (59%) and 9 of 34 isolates (26%) with geometric mean plasma titers between 1:40 and 1:125 (**Figure 6D, Table S6**). This was a substantial improvement in wild-type heterologous neutralization as compared to wild-type SHIV.CH505 infected RMs, whose plasma exhibited little to no heterologous neutralization.

**Figure 6.**
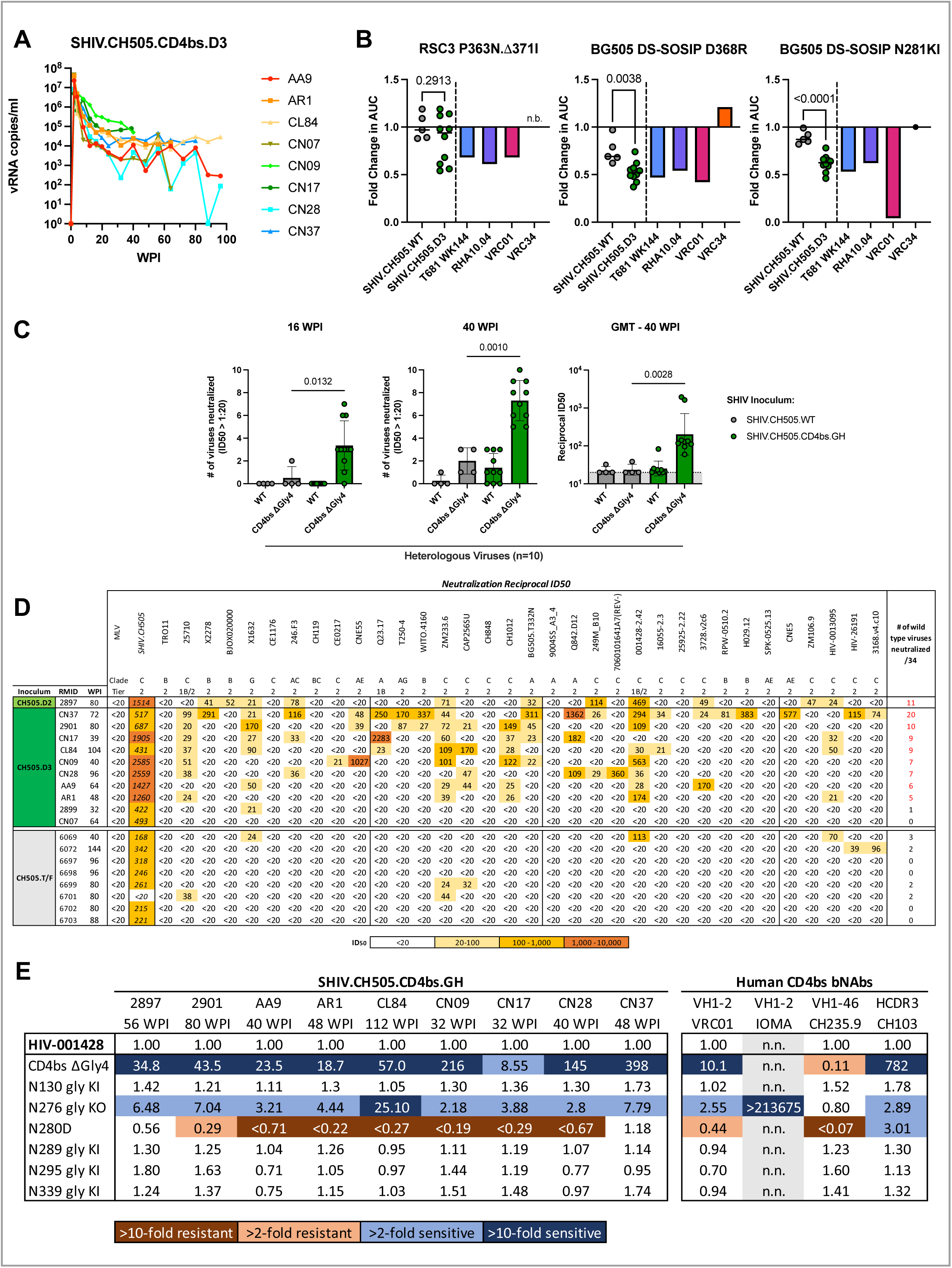
SHIV.CH505.CD4bs.GH elicited antibodies capable of broadly recognizing the underlying CD4bs backbone. (A) Viral load kinetics of eight additional SHIV.CH505.D3-infected macaques. (B) Fold change in binding AUC of rhesus plasma between WT and mutant RSC3 and BG505 DS-SOSIP proteins. Unpaired t-tests were used for comparing WT and GH groups. n.b, no binding to WT protein. (C) Neutralization (number and titer) of heterologous viruses with or without the CD4bs glycans by SHIV.CH505.WT and SHIV.CH505.CD4bs.GH-infected macaques. Error bars represent standard deviation from the mean. Significance determined by Mann-Whitney tests. GMT, geometric mean titer. (D) Heterologous plasma ID50 titers in SHIV.CH505.CD4bs.GH-infected macaques at peak breadth and in SHIV.CH505.WT infected macaques at euthanasia. (E) Neutralization ID50/IC50 fold change of SHIV.CH505.CD4bs.GH macaques with breadth and human CD4bs bNAbs against strain HIV-001428 and point mutations, colored by direction and magnitude. CD4bs βGly4, mutant lacks N197, N276, N363, and N462 glycans, gly KI, mutant restores a common M-group glycan, gly KO, mutant removes a common M-group glycan. N.n., not neutralized below 50 μg/mL.

To determine the epitopes targeted by the broadly neutralizing antibodies in these nine RMs, we evaluated changes in plasma neutralization titers against epitope-specific mutants of the heterologous strain HIV-001428-2.42 (**Figure 6E**). Knocking out N197, N276, N363, and N462 glycans together greatly increased neutralization titers in all nine RMs by 9 to 400-fold. Knocking out the N276 glycan alone also increased neutralization in all nine RMs by 2 to 25-fold. Introducing the Loop D mutant N280D reduced neutralization of HIV-001428 in seven of nine RMs by as much as 5-fold. Knocking in conserved glycans at residues 130, 289, 295, or 339 in HIV-001428 did not affect plasma neutralization titers from any of the SHIV.CH505.D3-infected macaque plasmas. Control human CD4bs bNAbs were affected as expected by these mutations^46–49^. Altogether, these results demonstrated that SHIV.CH505.CD4bs.D3 induced CD4bs targeted neutralization breadth in 8 of 10 infected macaques.

### Viral evolution in SHIV.CH505.CD4bs.GH-infected macaques revealed consistent sequential glycan restoration and CD4bs epitope escape

Viral epitope-specific sequence evolution provides a sensitive and specific indicator of the epitope specificity of NAb and CTL responses^4,7,49–54^. We performed longitudinal SGS on plasma vRNA from the ten SHIV.CH505.D3 infected and eight SHIV.CH505.T/F infected RMs^4^ (**Figure S6**). By 16 wpi, SHIV.CH505.D3 RMs had a significant net gain of PNGS across gp160 compared with SHIV.CH505.T/F animals (**Figure 7A**). These PNGS additions restored the engineered PNGS deletions in the SHIV inoculum and coincided temporally with a substantial rise in autologous neutralizing titers (**Tables S5, S6**). We further evaluated this glycan restoration by quantifying the change in glycan coverage of Env between the inoculum virus and the 16 wpi consensus sequences^55^; SHIV.CH505.D3 animals showed a significant mean increase of 1809 Å^2^ in glycan coverage by 16 wpi compared with control macaques (**Figure 7B**).

**Figure 7.**
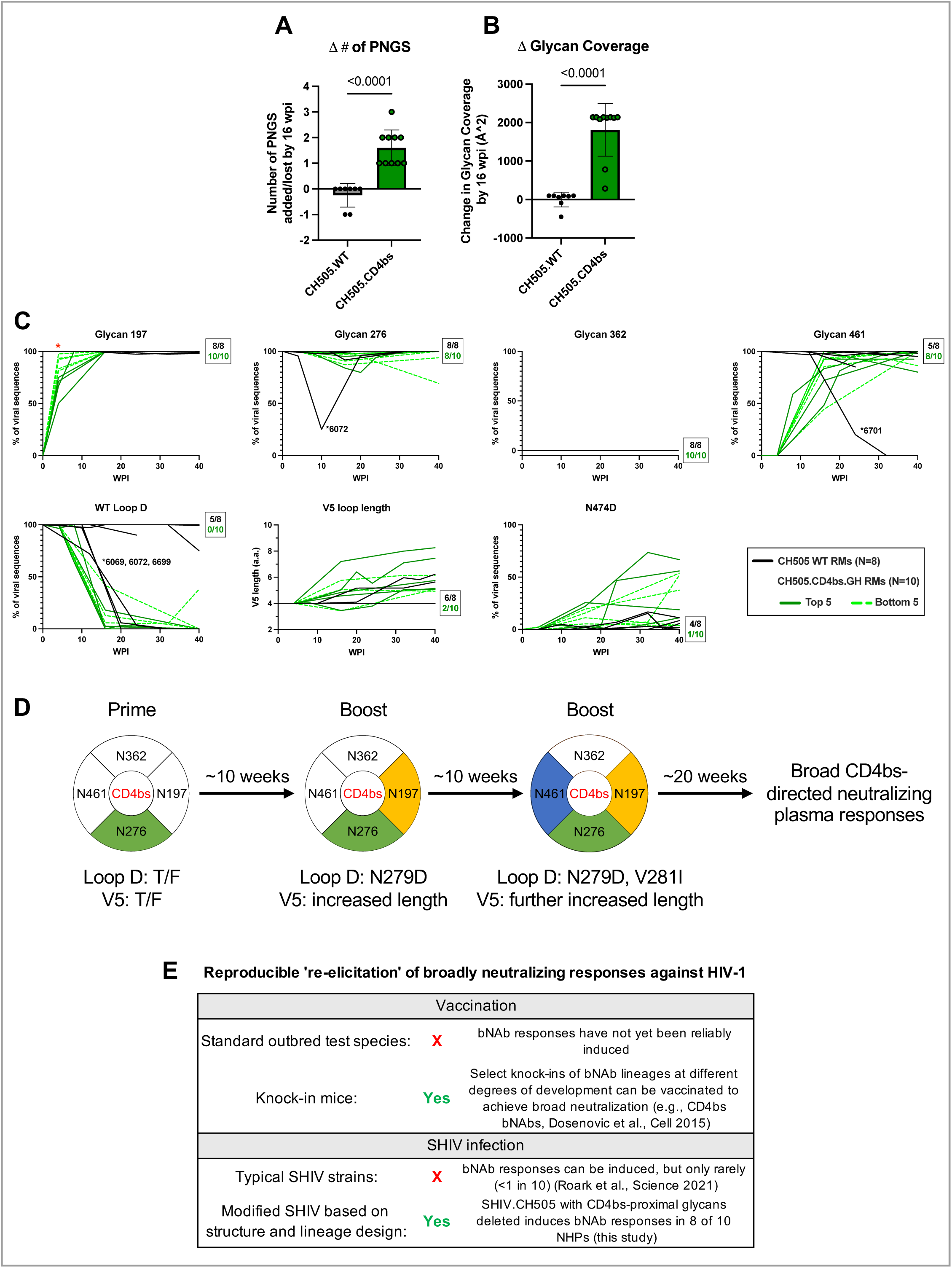
Sequential glycan reconstitution and CD4bs epitope viral evolution in SHIV.CH505.CD4bs.GH-infected macaques. (A) Comparison of the number of PNGS added by 16 wpi between WT and CH505.CD4bs.GH-infected macaques. Significance determined by Mann-Whitney Test. (B) Comparison of total glycan coverage of 16 wpi consensus sequences between WT and CH505.CD4bs.GH-infected macaques. Significance determined by Mann-Whitney Test. (C) SGS feature prevalence, thin lines represent individual macaques, thick lines represent group average. Boxed numbers represent the number of overlapping macaque lines at 40 wpi i.e. all 8 WT.SHIV.CH505 and all 10 SHIV.CH505.CD4bs.GH macaques lack a 362 PNGS in every Env sequence at 40 wpi. Solid green lines, Top 5 breadth SHIV.CH505.CD4bs.GH RMs. Dotted green lines, Bottom 5 SHIV.CH505.CD4bs.GH RMs. Mann-Whitney test applied to compare heterologous breadth in the top 5 and bottom 5 RMs, * = p<0.01 (D) Proposed immunization schema based on longitudinal sequencing in SHIV.CH505.D3 RMs. (E) Overview of CD4bs bNAb elicitation by vaccination or SHIV infection.

We next compared specific sequence changes longitudinally and between infection groups (**Figure 7C**). We observed a progressive restoration of the N197 and N462 glycans in SHIV.CH505.D3 RMs coinciding with mounting autologous neutralizing titers. Glycan 362, which is naturally absent in CH505.T/F, was not restored in any macaques in either group. The 276 glycan remained intact in all macaques, except for a control macaque RM 6702 where it underwent transient deletion at 10 wpi. Interestingly, a strain-specific CD4bs neutralizing mAb, DH650, was previously isolated from this macaque and was found to bind proximal to N276 in the autologous CH505.T/F Env^4^. Beyond PNGS mutations, we found known neutralization escape mutations at residues 279, 280, and 281 in Loop D in all ten SHIV.CH505.D3 RMs^46,56^. Variable loop 5 length variation is also known to be an escape pathway from CD4bs antibodies^49^, and we found that V5 lengthened in eight of ten SHIV.CH505.D3 RMs whereas six of eight SHIV.CH505.T/F RMs showed no change in V5 loop length. Finally, we observed a mutation N474D, which corresponds to a CD4-contact residue and was previously shown to represent an escape mutation from CD4bs bNAbs^46^, in nine of ten SHIV.CH505.D3 RMs. In three of these macaques, the N474D mutant was present in over half of the sequences by 40 wpi. In SHIV.CH505.T/F RMs, N474D was present in only half of the RMs and never above a quarter of the sequences.

We next asked if any longitudinal Env sequence features correlated with neutralization breadth development in SHIV.CH505.CD4bs.GH RMs. We split the macaques into two groups, high-breadth (N=5) and low-breadth (N=5), and evaluated the above features at each time point. The only significant feature that differentiated high-breadth RMs from low-breadth RMs was the proportion of sequences with the N197 PNGS restored by 4 wpi (**Figure S6B**). Higher breadth RMs had fewer sequences with an intact N197 PNGS than lower breadth RMs. We posit that in these RMs, the CD4bs glycan hole remained open longer than in lower breadth RMs, allowing for greater opportunity for suitable bNAb precursors to engage Env or more time for nascent bNAb lineages to affinity mature to accommodate wild-type glycans. Removing or underprocessing the N197 glycan has been explored successfully in CH235-like immunogen design^15^, and altogether, the data suggest that SHIVs with a higher threshold for restoring the N197 glycan may induce superior CD4bs antibody responses in infected macaques. Overall, the results with SHIV.CH505.D3 infection show how presentation of CD4bs-proximal glycan holes focuses B cell responses to the CD4bs followed by progressive filling of glycan holes and maturation of responses, eventually enabling >25% cross-clade neutralization breadth in 4 of 10 macaques and measurable cross-clade neutralization breadth in 8.

## DISCUSSION

The consistent induction of CD4bs-bNAbs has been a long-sought goal. In this study, we used clues from developmental analysis of a macaque CD4bs bNAb lineage to design modified SHIVs lacking CD4bs-proximal glycans as novel prime and boost “immunogens.” Infection of ten rhesus macaques by the glycan hole-variant SHIV.CH505.D3 induced broad HIV-1 neutralizing plasma responses in eight macaques, which was substantially higher than in 0 of 8 control macaques infected with SHIV.CH505.WT (p<0.001) or in 1 of 115 macaques infected with other wildtype SHIV strains (p<0.0001). Longitudinal plasma vRNA sequencing revealed sequential restoration of deleted glycans and CD4bs epitope escape, consistent with our hypothesis that infection by SHIV.CH505.D3 would immunofocus B cell responses to the CD4bs and be followed by Env-Ab coevolution, antibody affinity maturation, and acquisition of neutralization breadth including cross-clade responses.

Previous studies have suggested that RMs can serve as an important outbred animal model for the elicitation of HIV-1 bNAbs. This includes antibodies targeting the V2 apex^4,45,57^, V3 glycan^4,58–60^, MPER^61^, fusion peptide^54,62,63^, and CD4bs^4,8,9,15,64^. However, the consistent elicitation of broad and potent bNAb responses in the majority of infected or vaccinated macaques has remained elusive. The current study takes a significant step in this direction by showing that a structurally-guided, genetically-engineered SHIV depleted of glycans surrounding the CD4bs can elicit potent CD4bs-targeted autologous neutralizing antibody responses in 10 of 10 SHIV.CH505.D3 infected macaques and that in 8 of 10 macaques these responses could be affinity-matured to acquire neutralization breadth against at least 14% of a 34-member heterologous tier 2 virus screening panel. One macaque (2897) from the pilot study was infected with SHIV.CH505.D2 (**Figure 5C**), which lacked glycans at residues 362 and 461, and plasma from this macaque neutralized 11 of 34 (32%) of viruses (**Figure 6D**). Overall, plasma from 45% of macaques infected by CD4bs glycan hole-containing CH505 SHIVs neutralized >25% of heterologous viruses, with one (CN37) neutralizing nearly 60%. These results stand in stark contrast to 1 in 115 control wild-type SHIV infections that resulted in CD4bs bNAbs and is also quite different from the minimal neutralization breadth and potency observed by Sanders and Permar in BG505.GT1 SOSIP vaccinated macaques^8,9^, by Bjorkman and colleagues in SpyCatcher SOSIP nanoparticle vaccinated rhesus^64^, and by Saunders and colleagues in macaques vaccinated with glycan-modified CH505-based SOSIP nanoparticles^15^. These findings, together with the demonstration of broad (51%) and potent (0.436 µg/mL geomean IC_50_) neutralization by RHA10.04, suggest that the outbred rhesus model may be a valuable tool for HIV-1 vaccine design and development.

In macaque T681, we observed naturally occurring, transient CD4bs-proximal glycan deletions that preceded CD4bs bNAb elicitation, leading us to hypothesize that such deletions might enhance bNAb B cell precursor binding and activation. This is not the first association noted between spontaneous deletions in glycans in the evolving Env quasispecies and the subsequent priming of bNAb lineages^52,65^. In addition, we have observed spontaneous deletions at apical glycan residues N130, N160, or N187 leading to the development of V2 apex bNAbs in wildtype SHIV infected macaques^4,45,57^. Similarly, strains with glycan sparse V2 apexes are particularly sensitive to V2 apex bNAb precursors^66–68^, and SHIV infection with these strains frequently elicits V2 apex bNAbs^45^. Moreover, numerous prior studies^10,11,15,32–34,36,37,69,70^ have selectively deleted glycans surrounding the CD4bs, V3 high mannose patch, V2 apex, and fusion peptide epitopes as novel immunogen designs and have shown that this strategy consistently primes B cell responses to the respective epitope supersites. However, in none of these vaccination studies was it possible to boost responses to acquire consistent neutralization breadth and potency. The strategy that we developed here using glycan hole-containing SHIVs was specifically designed to address this requirement for boosting by providing a persistently replicating, diversifying, and coevolving immunogen. Nonetheless, since an infectious, replicating lentivirus can never be a vaccine, our long-range goal is to decipher SHIV Env-Ab coevolutionary pathways leading to neutralization breadth as a molecular guide or “blueprint” for prime and boost immunogen design (**Figure 7D**). Addressing this latter objective will be the focus of future studies.

Developing a safe and effective HIV-1 vaccine targeting the CD4bs remains a formidable challenge^12,71^ (**Figure 7E**). Two CD4bs antibody prototypes – VH gene-restricted CD4 mimetics and CDRH3-dominated – serve as principal targets for vaccine design, but it is not certain that a vaccine for either is achievable. The development of outbred nonhuman primate models for CD4bs bNAb induction can potentially aid and accelerate this effort and complement current promising human clinical trials^12^. The present study demonstrates unequivocally that rhesus monkeys can develop potent, broadly neutralizing CDRH3-dominated bNAbs, and the in-depth analysis of Env-Ab coevolution of the RHA10 bNAb lineage provides a detailed description of the intermediate steps in CH1012 Env evolution that were required. This included an unambiguous inference of the RHA10 lineage unmutated common ancestor (UCA) by a combination of IgDiscover and SONAR analyses and identification of a critical 4 a.a. insertion in the RHA10 lineage CDR-H3 that provided an affinity jump for evolving bNAb lineage B cells (**Figures 3A-C**). Importantly, Saunders recently reported that the rhesus IGHV1-105 gene encodes a CDRH2-dominated VH1-46-like CD4bs targeted bNAb-like precursor and that these precursors can be primed by glycan-modified CH505 nanoparticle immunogens^15^. It remains to be determined what CD4bs antibody type(s) are responsible for neutralization breadth in the other macaques of our present study and the developmental pathways by which they evolved. It is possible that by deciphering Env-Ab coevolution pathways in these macaques, boosting immunization strategies can be devised for both CDR-H2 and CDR-H3 dominated of CD4bs bNAb lineages.

A central theme in current HIV-1 vaccine research is that the consistent elicitation of bNAbs will require efficient priming of oftentimes rare naïve germline B cell precursors followed by efficient boosting and “polishing” of the expanding bNAb lineages to acquire neutralization breadth and potency while avoiding off-target responses. The design and testing of germline-targeted SHIVs in the rhesus model can facilitate and hasten the attainment of each of these objectives. SHIV infection has particular advantages as an immunization model. First, virus replication is high, persistent, and broadly disseminated throughout the lymphoreticular system, which equates to a high antigen load presentation to B cells^72,73^. Second, native Env trimers are expressed on the surfaces of virions and productively infected cells and they ensure correct presentation of canonical bNAb supersite epitopes^3,18^. Admittedly, non-native forms of Env are also expressed^74^ but epitopes on these proteins and polypeptides are likely to be largely different from immunodominant off-target responses such as those targeting the recombinant trimer base or glycan holes on native trimers that elicit strain-specific antibodies. In fact, the latter are often rapidly shielded in the evolving SHIV quasispecies by glycan additions^4,50,51,53,55^. Third, the evolving SHIV Env provides a continuous and rapidly changing affinity gradient or threshold to evolving B-cells in the expanding bNAb lineage because of its sensitivity to nascent neutralizing antibodies. Essentially, the evolving SHIV quasispecies rapidly escapes from nascent neutralizing antibodies, causing the affinity of all but a minor subset of the expanding bNAb lineage cells to drop and for these cells to apoptose and disappear, leaving only the rare cell(s) with higher affinity to persist and affinity-mature further. A prime example of this is illustrated here, where a single B cell that developed a 4 a.a. insertion in CDR-H3 acquired superior affinity for the evolving SHIV quasispecies and expanded at the expense of all other bNAb lineage B cells (see red lineage in **Figure 3B**). Fourth, because SHIV infection arises from a defined molecular viral clone and both Env and B cell evolution can be precisely and unambiguously inferred by single genome sequencing, next-generation deep-sequencing, and phylogenetic analyses, each step of bNAb induction can be deciphered and then further examined *ex vivo* and *in vivo* in iterative macaque infections or immunizations, or by immunizations in knockin mouse models. The current study is the first to illustrate many of these concepts, and related work is in progress with SHIVs that express germline-targeted Envs for other sites of vulnerability including V3 glycan, V2 apex, or fusion peptide epitope supersites^45,57,75^.

### Limitations of the study

While we demonstrate the reproducible induction of CD4bs-targeted antibodies with neutralization breadth, with the exception of the RHA10 lineage from T681, we have not isolated the corresponding bNAb mAbs. Isolation and characterization of such lineages will be necessary to determine the immunogenetic and structural classes of CD4bs bNAbs that SHIV.CH505.D2 and SHIV.CH505.D3 infection elicited and to define the Env-Ab coevolutionary pathways leading to neutralization breadth. These are high-priority goals for future research since such results will be critical for translating findings from the present study into novel prime and boost immunogen designs. A second limitation of the current study is that while the SHIV.CH505-glycan hole platform elicited heterologous neutralizing responses in RMs, the SHIV.BG505 and SHIV.CH1012 platforms did not. One potential explanation for this relates to the naturally missing CD4bs-proximal glycan at residue 362, which is missing from CH505 but not BG505 or CH1012, an explanation we are currently testing. A third limitation of the study is that the breadth and potency of CD4bs targeted neutralizing responses were, with two exceptions (RMs 2897 and CN37), modest compared with the RHA10 lineage and other human CD4bs bNAbs. This is not surprising, given that we prospectively induced bNAbs in a short period of time (<1 year) with our glycan hole-containing SHIVs, as opposed to retrospectively screening large numbers of chronically infected macaques or humans to identify those few individuals with high neutralization breadth and potency. Enhancing breadth and potency with new immunogen designs will be a priority going forward.

## Acknowledgments

Supported by the National Institutes of Health (GM103310, AI100148, AI160607, AI166916, AI016520, AI007632) and by the Vaccine Research Center, an intramural division of the National Institute of Allergy and Infectious Diseases, National Institutes of Health. D.J.M., R.S.R., and R.H. were supported by a Training Grant in HIV Pathogenesis (T32-AI007632). Some of this work was generated in the Penn Cytomics and Cell Sorting Shared Resource Laboratory at the University of Pennsylvania (P30-016520), the Simons Electron Microscopy Center (SF349247), and the National Resource for Automated Molecular Microscopy located at the New York Structural Biology Center (NYSTAR). We thank David Ambrozak at the Vaccine Research Institute for his aid in isolating RHA10.01 and the VRC Production Program for providing BG505 DS-SOSIP.664 Env trimer. The VRC Production Program includes Nadia Amharref, Nathan Barefoot, Christopher Barry, Elizabeth Carey, Ria Caringal, Kevin Carlton, Naga Chalamalsetty, Adam Charlton, Rajoshi Chaudhuri, Mingzhong Chen, Peifeng Chen, Nicole Cibelli, Jonathan W. Cooper, Hussain Dahodwala, Marianna Fleischman, Julia C. Frederick, Haley Fuller, Jason G. Gall, Isaac Godfroy, Daniel Gowetski, Krishana Gulla, Vera Ivleva, Lisa Kueltzo, Q. Paula Lei, Yile Li, Venkata Mangalampalli, Sarah O’Connell, Aakash Patel, Erwin Rosales-Zavala, Elizabeth Scheideman, Nicole A. Schneck, Zachary Schneiderman, Andrew Shaddeau, William Shadrick, Alison Vinitsky, Sara Witter, Yanhong Yang, and Yaqiu Zhang

## Author contributions

Conceptualization, D.J.M., P.D.K., G.M.S.; formal analysis, D.J.M., J.G., T.Z., A.J.C., H.L., R.H., C.A.S., K.W., B.T.K.; investigation, D.J.M., J.G., J.L., H.L., W.L., R.S.R., M.S.C., J.W.C., R.H., Y.L., C.L.M., Y.P., A.S., K.J.S, S.W., N.C., W.D., C.L., E.L., R.D.M., J.M.R.; resources, I.T., B.F.H.; writing – original draft, D.J.M, P.D.K., G.M.S.; writing – review and editing, all authors; supervision, B.T.K., M.S.S., D.C.D., B.F.H., D.W.K., M.R., B.H.H, P.D.K., G.M.S.; funding acquisition, B.F.H., P.D.K., G.M.S.

## STAR★ METHODS

### Key Resources Table

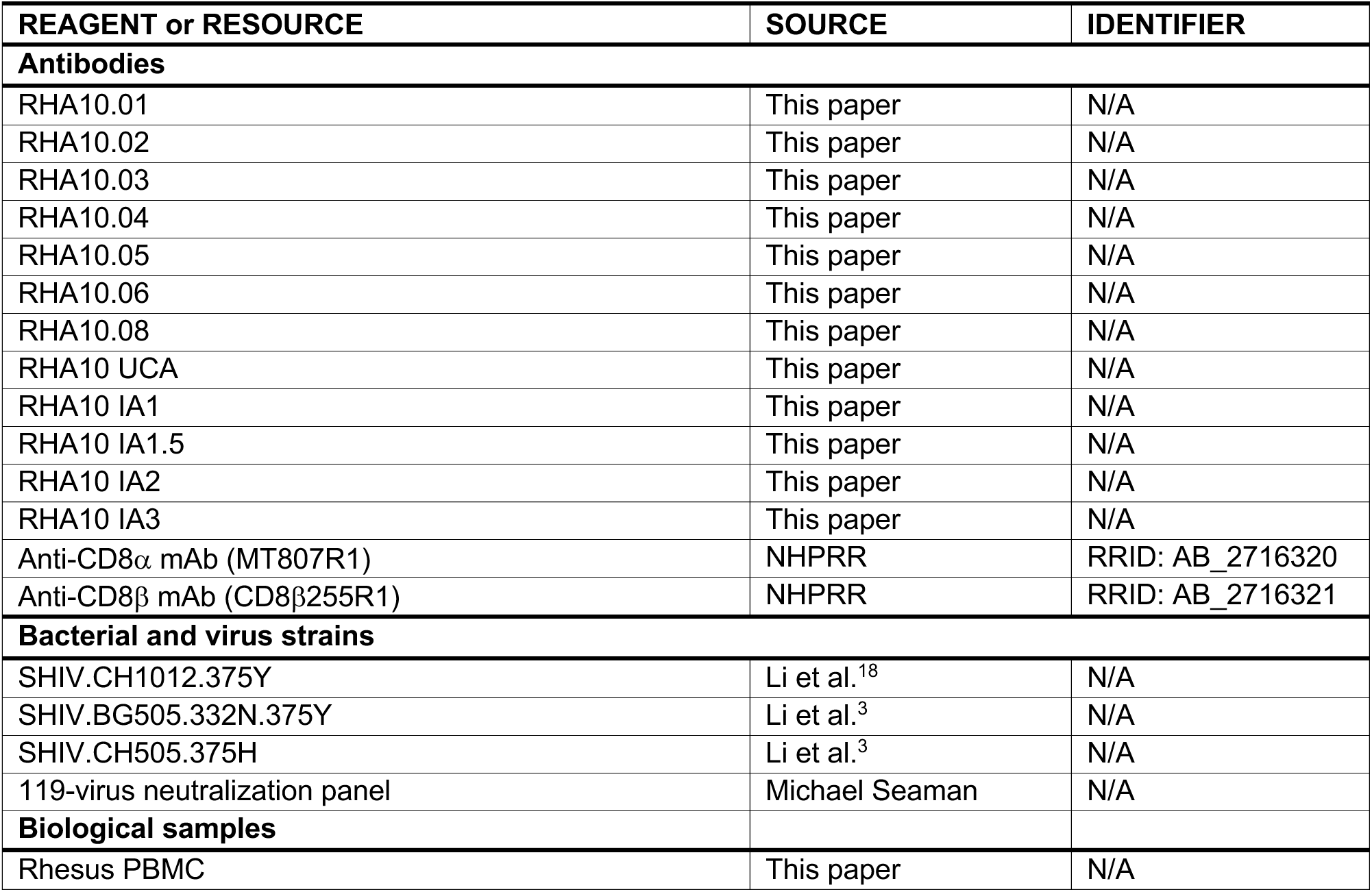
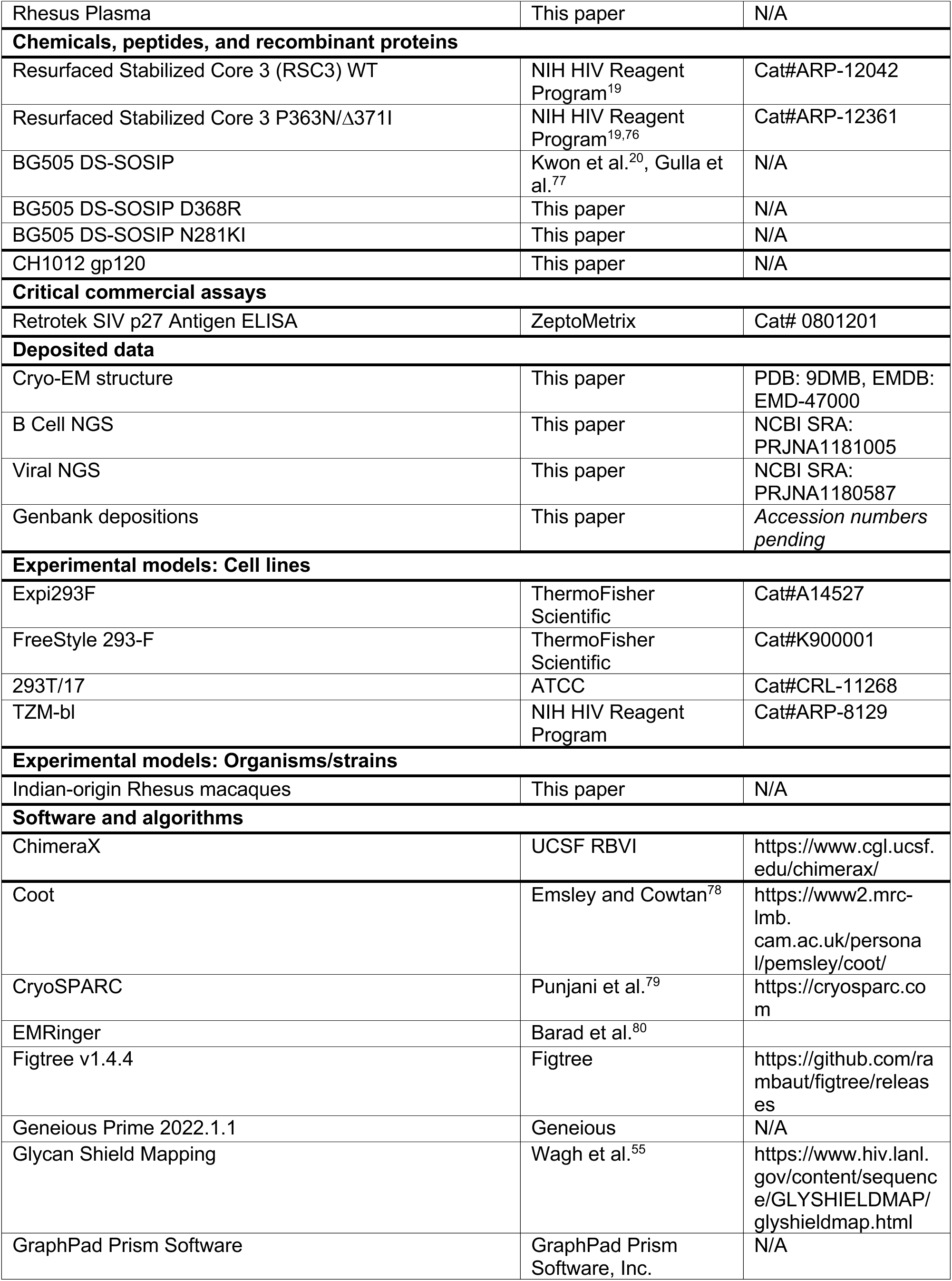
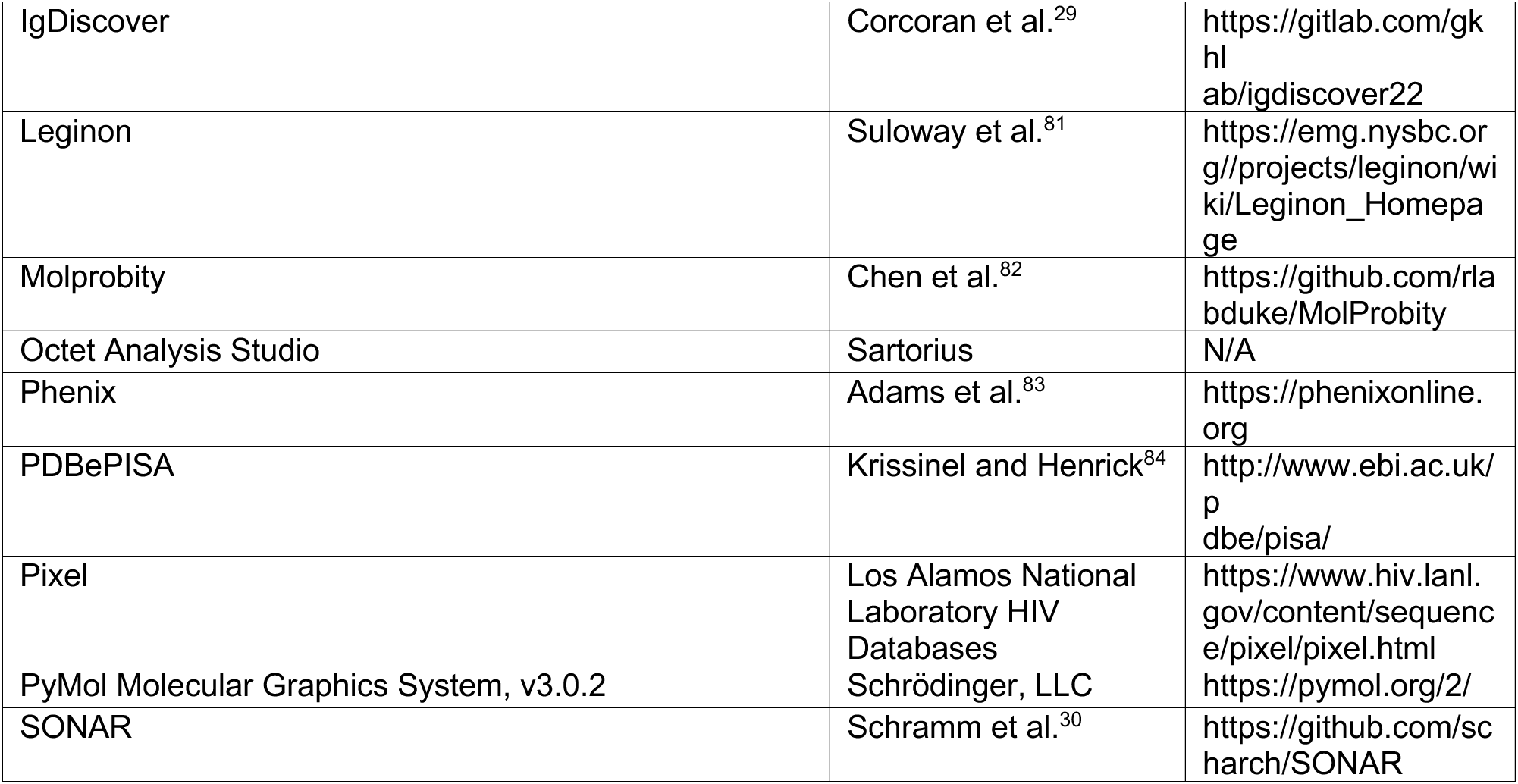

### Resource Availability

#### Lead Contact

Requests for further information should be directed to and will be fulfilled by George M. Shaw

#### Materials Availability

Requests for resources and reagents should be directed to and will be fulfilled by George M. Shaw. All new reagents are available by MTA for non-commerical research.

#### Data and Code Availability

GenBank accession numbers for RHA10 lineage members, UCA, and inferred intermediates are: PQ550578 – PQ550605. GenBank accession numbers for all HIV-1 *env* sequences analyzed in this study are as follows: CH1012 SGS, PQ574114 - PQ574448; CD4bs GH SGS, PQ574449 - PQ575767. GenBank accession numbers for IgDiscover-derived T681 immunoglobulin genes are: OR920055 – OR920194. GenBank accession numbers for CH1012 gp120 are: PP171710 – PP171713. Cryo-EM structure of RHA10.01 bound to BG505 DS-SOSIP is deposited under PDB ID 9DMB. Associated cryo-EM map was deposited to EMDB under EMD-47000. RHA10 lineage tracing NGS was deposited to NCBI SRA under PRJNA1181005. Viral NGS from macaque T681 was deposited to NCBI SRA under PRJNA1180587. Any additional information required to reanalyze the data reported in this paper is available from the lead contact upon request.

### Experimental Model and Study Participant Details

#### Nonhuman primates

Indian-origin rhesus macaques were housed at Bioqual, Inc., Rockville, MD, according to guidelines of the Association for Assessment and Accreditation of Laboratory Animal Care standards. Experiments were approved by the University of Pennsylvania and Bioqual Institutional Animal Care and Use Committees. RMs were sedated for blood draws, anti-CD8 mAb infusions, and SHIV inoculations.

#### Cell lines

293T/17 (ATCC #CRL-11268) , TZM-bl (NIH HIV Reagent Program, #ARP-8129), Expi293F (ThermoFisher Scientific Inc., #A14527), FreeStyle 293-F (ThermoFisher Scientific Inc, #R79007). These cell lines were cultured in line with manufacture suggestions as described in the below Method Details.

## Method Details

### SHIV infections

Previously, we reported the construction and *in vitro* and *in vivo* characterization of 16 novel SHIVs, each expressing a different primary HIV-1 Env^3,18^. We inoculated a total of 122 Indian-origin rhesus macaques with any one of these 16 SHIVs and followed them for 1-5 years for viral persistence and replication dynamics, the development of autologous and heterologous neutralizing antibodies, and clinical progression to AIDS-defining events. 7 macaques developed rapid clinical progression with persistently high plasma vRNA loads (>10^7^ vRNA molecules/ml of plasma) and failed to develop detectable anti-HIV Env or SIV Gag/Pol antibodies. These SHIV “rapid progressors” were similar to SIVmac239 rapid progressors^85^ and were euthanized <1 year after SHIV infection. Of the remaining monkeys, 115 were evaluable for the development of neutralization breadth^3,4,18,45^ (R.H. and G.M.S., *submitted*). The SHIV.CH1012 infected RMs T679, T680, T681, 6928, 6929 and 6930 analyzed in the current study were part of this large observational natural history study of SHIV infected animals. In the current study focused on Nab and bNAb development in SHIV.CH1012 infected RMs, a subset of these animals (6928, 6929, and 6930) received an intravenous infusion 25 mg/kg of anti-CD8beta mAb (CD8beta255R1) at the time of SHIV inoculation. The aim of anti-CD8 mAb treatment was to transiently deplete CD8+ cells, allowing for increased peak and setpoint viral loads. This was done in only a subset of animals because it was uncertain if this would accelerate disease progression to an extent that would preclude long term follow-up of animals for bNAb induction. In humans, higher plasma virus loads and lower CD4+ T cell counts are correlated with the development of bNAbs^16,86^. All SHIV infections were performed intravenously. The SHIV.CH1012.T/F inoculums contained molecularly cloned virus 300 ng p27Ag in RPMI1640 with filtered 10% heat-inactivated fetal bovine serum containing an equal mixture of Env375 variants. CD4bs glycan hole SHIV inoculums were the same but instead contained 100 ng p27Ag of molecularly cloned virus. All CD4bs.GH SHIV infected animals received an intravenous infusion of 25 mg/kg of anti-CD8alpha mAb (MT807R1) three days prior to SHIV inoculation.

### Processing and storage of clinical specimens

All blood samples were collected in sterile vacutainers containing ACD-A anticoagulant. 40 ml of ACD-A anticoagulated blood was combined in a sterile 50 ml polypropylene conical tube, centrifuged at 2100 rpm (1000xg) for 10 min at 20°C, and the plasma collected in a fresh 50 ml conical tube, avoiding disruption of the buffy coat WBC layer and large red blood cell pellet. The plasma was centrifuged again at 2500 rpm (∼1500g) for 15 min at 20°C in order to remove all platelets and cells. Plasma was collected and aliquoted into 1 ml cryo-vials and stored at −80°C. The RBC/WBC pellet was resuspended in an equal volume of Hanks balanced salt solution (HBSS) without Ca++ or Mg++ and containing 2mM EDTA and then divided into four 50 ml conical tubes. Additional HBSS-EDTA (2 mM) buffer was added to adjust the volume of the RBC/WBC mixture to 30 ml in each tube. The cell suspension was then gently underlayered with 14 ml 96% Ficoll-Paque and centrifuged at 1800 rpm (725xg) for 20 min at 20°C in a swinging bucket tabletop centrifuge with slow acceleration and braking to maintain the ficoll-cell interface. Mononuclear cells at the ficoll interface were collected and transferred to a new 50ml centrifuge tube containing HBSS-EDTA (w/o Ca++ or Mg++) and centrifuged at 1000 rpm (∼200g) for 15 min at 20°C. This pellets PBMCs and leaves most of the platelets in the supernatant. The supernatant was discarded, and the cell pellet was resuspended in 40 ml HBSS (with Mg++/Ca++ and without EDTA) + 1% FBS. To further remove contaminating platelets, the cell suspension was centrifuged again at 1000 rpm (∼200g) for 15 min at 20°C and the supernatant discarded. The cell pellet was tap-resuspended in the residual 0.1 to 0.3 ml of media and then brought to a volume of 10 ml HBSS (with Mg++/Ca++) + 1% FBS. Cells were counted and viability assessed by trypan blue exclusion. Cells were centrifuged again at 1200 rpm (300xg) for 10 min at 20°C, the supernatant discarded, and the cells resuspended at a concentration of 5 to 10 × 10^6 cells/ml in CryoStor cell cryopreservation media (Sigma Cat. C2999) and aliquoted into 1-ml cryovials (CryoClear cryovials; Globe Scientific Inc., Cat. 3010). Cells were stored in a Mr. Frosty at −80°C overnight and then transferred to vapor phase liquid N2 for long-term storage.

### SHIV construction and characterization

The experimental design for constructing SHIVs bearing primary or transmitted/founder Envs with allelic variation at gp120 residue 375 was previously described, including SHIV.CH1012^3^. CD4bs Glycan Hole SHIVs were constructed from the original base SHIV strain and PNGS mutations were made using Q5 Site-Directed Mutagenesis Kit (NEB). Each viral genome was sequenced in its entirety to authenticate its identity. Infectious SHIV stocks were generated in 293T cells as previously described^3^. On day 0, five million 293T cells were plated in 100-mm tissue culture dishes in 10 ml of complete DMEM growth media with 10% FBS. On day 1, 6 ug of SHIV plasmid DNA was combined with 18 uL of FuGENE 6 (Promega) in 500 μl of DMEM was added dropwise to tissue culture dishes. Media containing virus was harvested on day 3 and aliquoted for storage at −80°C. Virus concentration was estimated by p27 antigen (p27Ag) ELISA (Zeptometrix) and infectious particle concentration was determined by entry into TZM-bl cells in the presence of DEAE–Dextran, as previously described^50^. The replication kinetics of each of the SHIV.CH1012 Env375 variants in primary, activated human and rhesus CD4 T cells were determined as previously described ^3^. 293T supernatants containing 300 ng p27Ag of each variant, were added to 2 × 10^6 purified human or rhesus CD4 T cells in complete RPMI growth medium (RPMI1640 with 15% FBS (Hyclone), 100 U/ml penicillin–streptomycin (Gibco), 30 U/ml IL-2 (aldesleukin, Prometheus Laboratories) and 30 μg/ml DEAE-Dextran. The cell and virus mixtures were incubated for 2 hours under constant rotation at 37C to facilitate infection, washed three times with RPMI1640, and resuspended in complete RPMI1640 medium lacking DEAE-Dextran. Cells were plated into 24-well plates at 2 million cells in 1 ml and cultured for 11 days, with sampling of 0.2 ml supernatant and media replacement every 2 to 3 days for 11 days. Supernatants were assayed for p27Ag concentration by ELISA (Zeptometrix).

### Plasma vRNA quantification

Plasma viral load measurements were performed by the NIH/NIAID-sponsored Nonhuman Primate Virology Core Laboratory at the Duke Human Vaccine Institute. This core facility is CLIA certified and operates a highly standardized, quality-controlled Applied Biosystems Real-Time SIV and HIV vRNA PCR assays. QIAsymphony SP and QIAgility automated platforms (QIAGEN) were used for high throughput sample processing and PCR setup. Viral RNA was extracted and purified from plasma, annealed to a target specific primer and reverse transcribed into cDNA. The cDNA was treated with RNase and added to a custom real-time PCR master mix containing target specific primers and a fluorescently labeled hydrolysis probe. Thermal cycling was performed on a QuantStudio3 (ThermoFisher Scientific) real-time quantitative PCR (qPCR) instrument. Viral RNA cp/reaction was interpolated using quantification cycle data. Raw data were quality-controlled, positive and negative controls are checked, and the mean viral RNA cp/ml was calculated. The lower limit of quantification (LLOQ) of this assay was 62 RNA copies/mL.

### Neutralizing antibody assay

Assays for neutralizing antibodies were performed using TZM-bl indicator cells, as previously described^3,50^. This assay is essentially identical to that employed by Montefiori, Seaman and colleagues^87^ (www.hiv.lanl.gov/content/nab-reference-strains/html/home.htm), the only difference being that in our assay we plate virus and test plasma onto adherent TZM-bl cells and hold the concentration of test plasma constant (5% vol/vol) constant across all wells, which contain 10% heat-inactivated normal human serum (Gibco) in the complete DMEM culture medium. Target viruses express HIV-1 Envs whose complete designations and subtype classifications are reported elsewhere^88,89^.

### Enzyme-linked immuosorbent assays (ELISAs)

#### RSC3, CH1012 gp120s

Half-area 96-well (Corning, Cat. #3690) plates were coated overnight at 4°C with anti-His tag mAb (Thermofisher, Cat. #PA1-983B) then washed three times with 0.2% (v/v) Tween-20 supplemented phosphate buffered saline (PBST). Plates were then blocked for 1 hour at room temperature with 5% (w/v) fat-free milk powder dissolved in PBST and the blocking buffer was shaken out by hand. 0.25 μg of recombinant gp120 subunits diluted in PBS were added to the wells for 1 hour at room temperature and washed as above. 4-fold serially diluted mAbs were added to wells in duplicate for 1 hour at room temperature and washed as above. Antibody binding was detected with Peroxidase-conjugated AffiniPure Donkey Anti-Human IgG (H+L) (Jackson ImmunoResearch Laboratories, RRID: AB_2340495) followed by a PBST wash. HRP detection was quantified with addition of 3,3′,5,5′-tetramethylbenzidine (TMB) (Sigma Aldrich, Cat. # T0440) for 10 mins and this reaction was stopped by the addition of 1 N sulfuric acid (Ricca, Cat. #8300-32). HRP activity was determined as the optical density at 450 nm subtracted by the background optical density at 570 nm.

#### BG505 DS-SOSIPs

Plasma and mAb binding to BG505 DS-SOSIP and mutants were performed as above but the 96-well plates were coated instead with PGT128 Fab at 2 µg/mL.

### Rhesus B cell isolation

RHA10.01 was isolated from cryopreserved PBMC from week 72 post-SHIV infection as previously described^4^. Briefly, cells were thawed and stained with LIVE/DEAD Fixable Aqua Dead Cell Stain (Life Technologies), as previously described^90,91^. Cells were washed and stained with an antibody cocktail of CD3 (clone SP34-2, BD Biosciences), CD4 (clone OKT4, BioLegend), CD8 (clone RPA-T8, BioLegend), CD14 (clone M5E2, BioLegend), CD19 (clone CB19, Novus Bio.), CD20 (clone 2H7, BioLegend), IgG (clone G18-145, BD Biosciences), IgD (polyclonal, Dako), IgK (clon SPM558, Novus Bio.), IgL (MHL-38, BioLegend), and biotinylated rhesus IgA (clone 10F12, NHPRR) at room temperature in the dark for 20 min. The cells were then stained with a streptavidin-Brilliant Violet 421 for 15 min at room temperature. The stained cells were washed 3 times with PBS, resuspended in 1 ml of PBS and passed through a 70 μm cell mesh (BD Biosciences). Total memory B cells (CD3-CD4-CD8-CD14-, CD19 or CD20+, IgD-IgM-IgA-IgG+, IgK or IgL+) were sorted with a modified 3-laser FACSAria cell sorter using the FACSDiva software (BD Biosciences) and flow cytometric data was subsequently analyzed using FlowJo (v10.10). B cells were sorted at 1 cell per well of a 384-well plate containing B cell culture media based on a human B cell culture protocol^92^ that was optimized for the expansion of rhesus B cells. Briefly, sorted B cells were expanded for 14 days in B cell culture medium consisting of Iscove’s modified Dulbecco’s medium (IMDM) with GlutaMAX supplemented with 10% heat-inactivated fetal bovine serum (FBS), 1X MycoZap Plus-PR, 100 U/ml IL-2, 0.05 ug/ml IL-4, 0.05 ug/ml IL-21, 0.05 ug/ml BAFF, 2 ug/ml CpG ODN2006, and 3T3-msCD40L feeder cells at a density of 5000 cells per well. Supernatants from ∼20,000 individual wells were evaluated for neutralization of HIV-1 BG505.T332N and X1632_S2_10 Env-pseudotyped viruses using the high throughput NVITAL automated micro-neutralization assay, as previously described^93^. RHA10.02 through RHA10.08 were isolated from 144 wpi PBMCs and were similarly stained as above. These cells were then sorted into 96-well plates containing lysis solution and proceeded directly to single-cell PCR described below. Wells with amplified RHA10 sequences were cloned and expressed as below.

### Rhesus B cell cloning and expression

Heavy (IGHV) and light (IGKV, IGLV) chain genes were isolated via single-cell PCR approaches^94,95^. For mAb isolation from RM T681, bulk cDNA was synthesized from the sixteen neutralization-positive B cell culture wells using random hexamers as previously described^96^. Subsequently, immunoglobulin heavy chain (IgG) and light chain (IgK and IgL) variable regions were separately amplified by nested PCR cycles using pools of rhesus macaque primers as previously described^21^. The heavy and kappa chain variable regions of the RM T681 lineage were codon optimized, synthesized with a murine immunoglobulin signal peptide (tripeptide sequence VHS) immediately preceding the 5′ end of FRW1, and cloned into rhesus IgG1 (RhCMV-H) and IgK (RhCMV-K) expression vectors directly upstream of the respective constant regions using AgeI/NheI and AgeI/ScaI restriction sites, respectively^21^. Recombinant mAbs were expressed by cotransfection of paired heavy and lambda chain plasmids into 293Freestyle cells as previously described^19^, purified from cell supernatant using the Protein A/Protein G GraviTrap kit (Cytiva) and buffer-exchanged into PBS.

### Neutralization correlation analyses

Hierarchical clustering using Log10 IC50 titers was performed using heatmap webtool on the LANL HIV Databases (hiv.lanl.gov). Cluster robustness was assessed using 100 bootstraps and only those splits with >50% are indicated in Figure 1F. Neutralizing Log10 IC50 titers for RHA10.04 versus other bNAbs were plotted together and Pearson correlation test was used to identify significantly correlated titers. Fisher’s exact test was used to assess significance of the overlap in simultaneously sensitive or simultaneously resistant pseudoviruses between RHA10.04 and other bNAbs. To explore hypotheses, all uncorrected p < 0.05 associations were deemed interesting, but given that 38 tests were done, only the tests with uncorrected p < 0.001 would be significant after a Bonferroni correction for multiple tests.

### Soluble HIV-1 envelope trimer generation

HIV Env trimer BG505 DS-SOSIP.664 was produced in transiently transfected 293F cells as previously described^97,98^. Briefly, the plasmid encoding the SOSIP trimer and a plasmid encoding furin were mixed at 4:1 ratio and transfected into 293F cells at 0.75 mg plasmid per 1 L cells (1.5 × 10^6/ml) using 293Fectin (ThermoFisher) or Turbo293 transfection reagent (Speed BioSystems). Cells were incubated in a shaker at 120 rpm, 37°C, 9% CO2. The following day, 80 ml HyClone SFM4HEK293 medium and 20 ml Free-Style 293 Expression Medium were added to each liter of cells. Env trimer protein was purified from the day-7 super-natant with VRC01 affinity chromatography, followed by gel filtration on a Sephadex200 16/60HL column.

### Antibody Fab preparation

Variable regions of the RHA10.01 antibody heavy and light chain genes were synthesized (Genscript) and each subcloned into the pVRC8400 vectors, in which a HRV3C cleavage site was inserted in the heavy-chain hinge region. The heavy and light chain pair was co-transfected in Expi293F cells (ThermoFisher) using Turbo293 transfection reagent (Speed Bio-Systems) as described previously^99^. The culture supernatant was harvested 6 days post transfection and loaded on a protein A column, the column was washed with PBS, and IgG proteins were eluted with a low pH glycine buffer and the pH was immediately neutralized with Tris pH 8. The eluted IgG proteins were cleaved by HRV3C, and the cleavage mixture was passed through a protein A column.

### Cryo-EM data collection and processing

Antibody Fab fragments of RHA10.01 were incubated with BG505 DS-SOSIP with the Fab in two fold molar excess per protomer. 2.3 μl of the complex at 1 mg/ml concentration was deposited on a C-flat grid (www.protochips.com). An FEI Vitrobot Mark IV was used to vitrify the grid with a wait time of 30 s, blot time of 3 s and blot force of 1. Automated data collection was performed with Leginon^81^ on a Titan Krios electron microscope equipped with a Gatan K3 Summit direct detection device. Exposures were taken in movie mode for 5 s with the total dose of 70.72 e–/Å2 fractionated over 50 raw frames. Images were pre-processed through Appion^100,101^; MotionCor2^102^ was used for frame alignment and dose-weighting. The CTF was estimated using CTFFind4^103,104^. Initial particle picking was done with DoG Picker^100,101^. RELION^105^ was then used for particle extraction. CryoSPARC 2.12^79^ was used for 2D classifications, ab initio 3D reconstruction, homogeneous refinement, and nonuniform 3D refinement. Initial 3D reconstruction was performed using C1 symmetry, confirming 3 Fabs per trimer and C3 symmetry was applied for the final reconstruction and refinement. Simulated annealing was performed on the first refinement followed by iterative manual fitting and real space refinement in Phenix^83^ and Coot^78^. Geometry and map fitting were evaluated throughout the process using Molprobity^106^ and EMRinger^80^. PyMOL (www.pymol.org) was used to generate figures. Buried Surface Area (BSA) between the RHA10.01 heavy chain and BG505 DS-SOSIP was calculated using the PISA server^84^.

### Angle of approach comparisons

The angles of approach for CD4 and CD4-binding site antibodies on the trimeric Env were defined as described previously^5^. In brief, the trimer coordinate system is defined by the trimer 3-fold axis and a perpendicular axis passing the Cα atom of residue Asp368 in of the Env protomers. The latitudinal angle measures the freedom of movement between the viral membrane with a latitudinal angle of 0 coinciding with the trimer 3-fold axis. The longitudinal angle measures the freedom of movement between neighboring Env protomers. The axis of antibody Fab is defined by two points where the coordinates of one point are the average of Cα atoms of Cys residues in both heavy chain VH and light chain VL domains, and the coordinates of the second point are the average of Cα atoms of Cys residues in both heavy chain CH1 and light chain CL domains.

### Rhesus B Cell Lineage Tracing

#### Individual rhesus antibody allele repertoire

PBMCs isolated from RM T681 plasma obtained at week 72 post-infection were stained with LIVE/DEAD Aqua, CD3-PerCP-Cy55, CD4-BV785, CD8-BV711, CD14-PE-Cy7, CD20-BV605, IgD-FITC, IgG-AF700, and IgM-BV650. Approximately 50,000 naive B cells (CD20+, IgG−, IgD+, IgM+) were bulk sorted into RPMI with 10% FBS and 1% Pen-Strep using a BD FACSAria II. Total RNA was extracted using RNAzol RT per the manufacturer’s guidelines (Molecular Research Center, Inc). Reverse transcription of mRNA transcripts, IgM and IgK variable region library preparation, and next generation sequencing were performed as previously described^107^, as these methods can be efficiently used for both humans and rhesus macaques. Both heavy and kappa immunoglobulin libraries were sequenced on Illumina Miseq with 2×300 bp runs using the MiSeq Reagent V3 kit (600-cycle). Filtered, quality-controlled IgM and IgK sequences were analyzed using IgDiscover^29^ to curate a personalized immunoglobulin repertoire library for RM T681. A naïve IgL variable region library was not prepared since the RHA10 utilizes a kappa chain. IgDiscover can identify functional VH, VL, VK, JH, JL and JK genes, and denotes any novel genes and/or alleles with respect to the provided reference database. For this analysis, a recently published rhesus macaque database was used as the template^108^.

#### IgG+ RHA10 lineage tracing

We used SONAR^30^ for longitudinal analysis of antibody sequences from IgG+ sorted PBMCs, starting with annotation and clustering, proceeding to manually-guided candidate lineage member selection using the identity/divergence plot feature, and ending with phylogenetic analysis across timepoints with SONAR’s IgPhyML-based features. In addition to the phylogenetic tree for each chain, SONAR produces a set of inferred ancestor sequences for the internal tree nodes spanning the UCA to the mature antibody sequences. We selected ancestor sequences for the heavy and light chains to pair together as functional intermediate antibodies. Sequences were selected to provide a representative series of on-track phases of somatic hypermutation over time toward the mature bNAb lineage members, both before and after the CDR-H3 insertion. We sought to match, as best as possible given the unpaired nature of the underlying data, similarly-progressed heavy and light chains in terms of levels of SHM and the context of observed members by timepoint from each tree.

### Viral Sequencing and Visualization

#### Single genome sequencing (SGS)

SGS of SHIV 3′ half genomes was performed as previously described^2,3^. Geneious Prime 2022.1.1 (https://www.geneious.com) was used for alignments and sequence analysis and sequences were visualized using the LANL *Pixel* tool^109^ (https://www.hiv.lanl.gov/content/sequence/HIV/HIVTools.html). Phylogenetic trees were generated and visualized using Geneious Prime 2022.1.1 and Figtree v1.4.4. The specific implementation of this software for this project is described in the figure legends.

#### Viral NGS

Approximately 100,000 viral RNA copies were extracted from plasma virions by using the Qiagen BioRobot EZ1 Workstation with EZ1 Virus Mini Kit v2.0. vRNA was eluted and immediately subjected to cDNA synthesis. Reverse transcription of RNA was performed by using MuLV (SuperScript III) reverse transcriptase and SIVmac251.R.R1 reverse primer according to manufacturer’s instructions. To prevent resampling the PCR products, we obtained PCR end point dilution of each cDNA sample by using single genome amplification method and calculated the actual number of molecules amplified per μl of cDNA. Based on this result, we determined the volume of cDNA used for population PCR amplification to reach a sampling depth of at least >2,500 individual molecules. We ran multiple population bulk PCR reactions and then pooled the PCR product together for MiSeq analysis if the cDNA volume needed was too large to fit into one PCR. Primers used to amplify Env regions contain both 362 and 461 PNGS were as follows: CH1012.RHA10.F1 5’-GCATGGAATAAAACTTTACAAGCGGT-3’ (nt 7,233 – 7,257, HXB2), CH1012.RHA10.R1 5’-CCACTTTATATTTATATAATTCACTTCTC-3’ (nt 7,661 – 7,689, HXB2). PCR products were run on a 1% agarose gel and the correctly sized DNA band was purified using PureLink Quick Gel Extraction kit (Invitrogen). Samples were then sequenced using an Illumina MiSeq 500V2 Nano kit using a 20% PhiX spike-in. High-quality reads were trimmed, paired, merged, deduplicated, and aligned to the CH1012 Env reference sequence. Reads that assembled to the Env and spanned both PNGS were compiled and analyzed by a custom script to analyze the reads for presence or absence of the PNGS of interest.

### Recombinant protein expression and purification

In order to produce CH1012 gp120s, an L111A mutation was introduced into sequences to prevent dimerization^110^. Gp120 DNA was synthesized and cloned into a pVax expression plasmid (Genscript). The proteins were produced by transfection of 293F cell cultures in Expi293 expression media (ThermoFisher) using Expifectamine transfection reagent (ThermoFisher) according to the manufacturer’s instructions. The protein-containing supernatant was harvested by centrifugation at 3000g for 30 minutes, followed by filtration of the supernatant (0.22µm Stericup Quick Release filter, Millipore Sigma). Gp120s were purified from the filtered supernatant by lectin-affinity chromatography using lectin beads (Vector Laboratories), eluted in 1M Methyl alpha-D mannopyranoside, and further purified by size-exclusion chromatography in PBS (GE S200 increase column).

### Biolayer interferometry

Binding KDs were determined using an Octet R8 protein analysis system (Sartorius). RHA10.04 or RHA10 UCA IgG was diluted in PBST with 1% BSA to a concentration of 5µg/ml and captured on anti-human IgG-Fc biosensors (Sartorius) for 300s. Association was measured by incubating the loaded sensors with gp120s for 300s, and dissociation was measured by incubating the sensors in PBST with 1% BSA for 600s. For gp120s, a starting concentration of 2µM followed by eight 3-fold dilutions was used for RHA10.04 affinity measurements, and a starting concentration of 50µM followed by four 2-fold dilutions was used for RHA10 UCA affinity measurements. Data was analyzed using Octet Analysis Studio Software (Sartorius).

### Statistical analyses

Statistical tests were performed using GraphPad Prism 10 software. The specific usage is described in the figure legends.

## Summary of Supplemental Information

**Figure S1.**
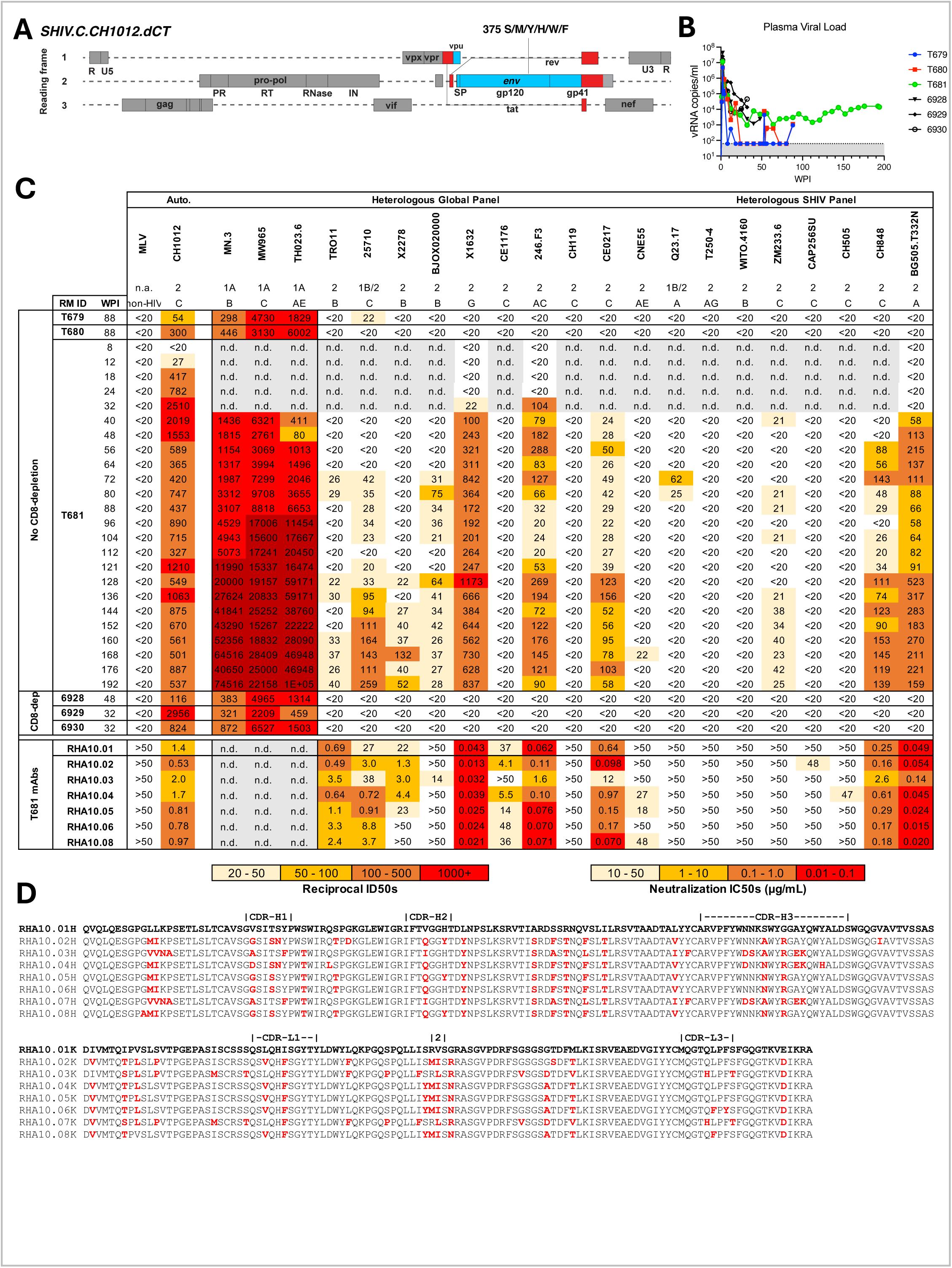
SHIV.CH1012 is infectious in rhesus macaques and elicited heterologous neutralization in macaque T681 that is recapitulated by the RHA10 lineage. **A.** SHIV genetic organization schematic displaying the gene organization and strain contribution to the final virus. Gray – SIVmac766, Red – HIV.D.191859, Blue – HIV.C.CH1012 T/F. **B.** Longitudinal plasma viral loads of SHIV.CH1012-infected macaques. RMs 6928, 6929, and 6930 were treated with anti-CD8α three days prior to infection. RMs T679 and T680 we treated with anti-CD8α at 52 WPI. Lower limit of quantification = 62 copies/mL. **C.** SHIV.CH1012-infected macaque longitudinal plasmas and RHA10 members were tested for neutralization against murine leukemia virus (MLV), autologous SHIV.CH1012, Tier 1A viruses, heterologous Global Env Diversity Panel, and heterologous Tier 2 SHIV panel. Values represented as reciprocal inhibitory dilution 50 (ID50s). n.d. = not determined. Lower limit of detection = 20 for plasmas, 50 ug/mL for mAbs. RHA10.02 and RHA10.07 are identical at the amino acid level therefore only RHA10.02 was screened for neutralization **D.** Alignments of RHA10 lineage members heavy and kappa chain variable regions according to IMGT numbering. Disagreements with the RHA10.01 heavy and kappa chains are highlighted. RHA10 CDR-L2 is 3 amino acids long and is labeled as “2” in the alignment. GenBank accession numbers for RHA10 heavy and kappa chains are: PQ550578 – PQ550585, PQ550592 – PQ550599.

**Figure S2.**
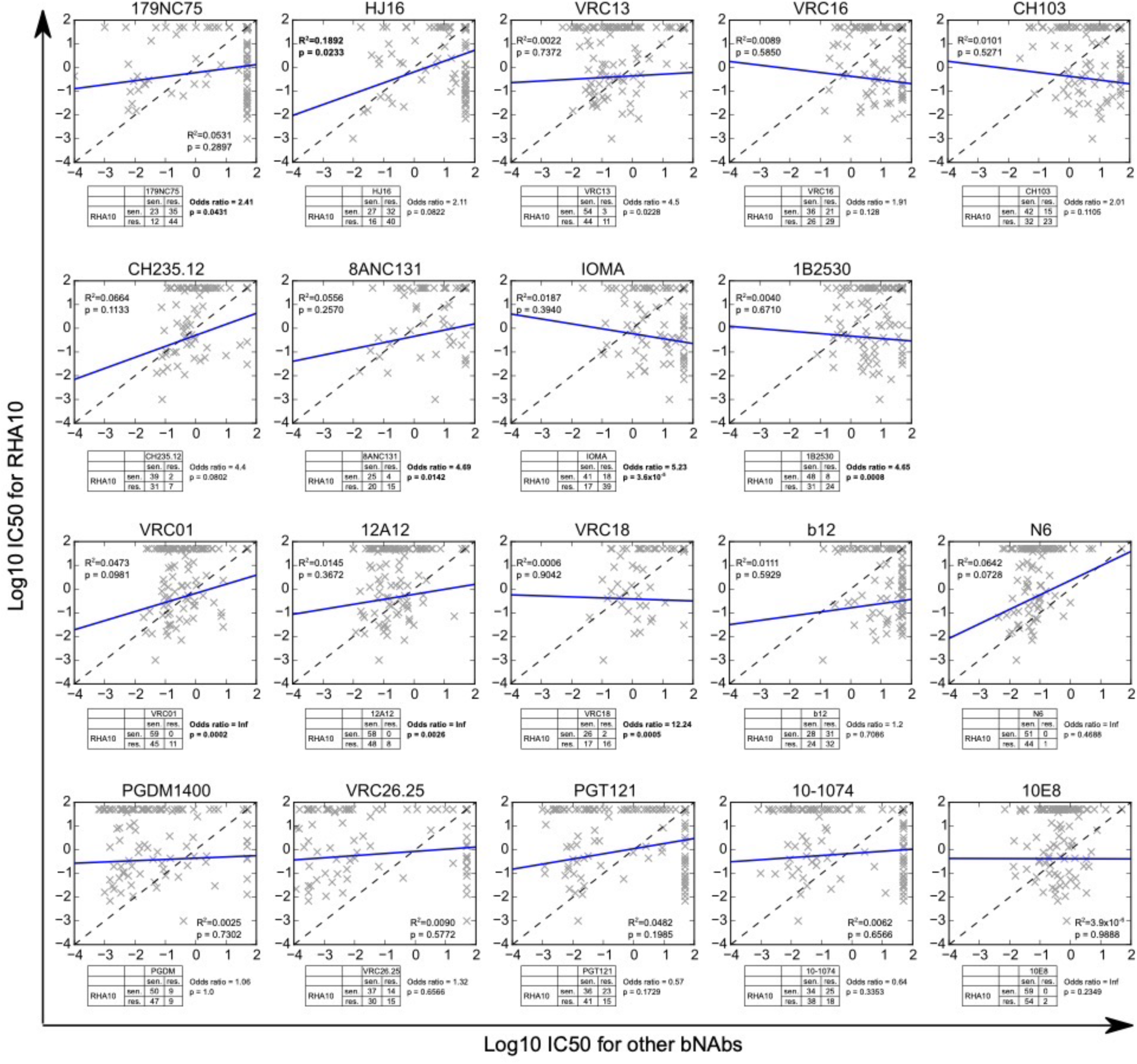
Correlation analyses of RHA10 neutralizing titers. Each panel shows correlation between Log10 IC50 titers for RHA10 on the vertical axis, and for each comparator bNAb on the horizontal axis. Each point is titers against same heterologous pseudovirus. R2 and p-values from Pearson correlation test are shown in the plot. The table below each plot shows the number of heterologous viruses sensitive/resistant for RHA10 and each comparator bNAb, and odds ratio and p-value from Fisher’s exact test are shown. Any tests with uncorrected p < 0.05 are shown in bold

**Figure S3.**
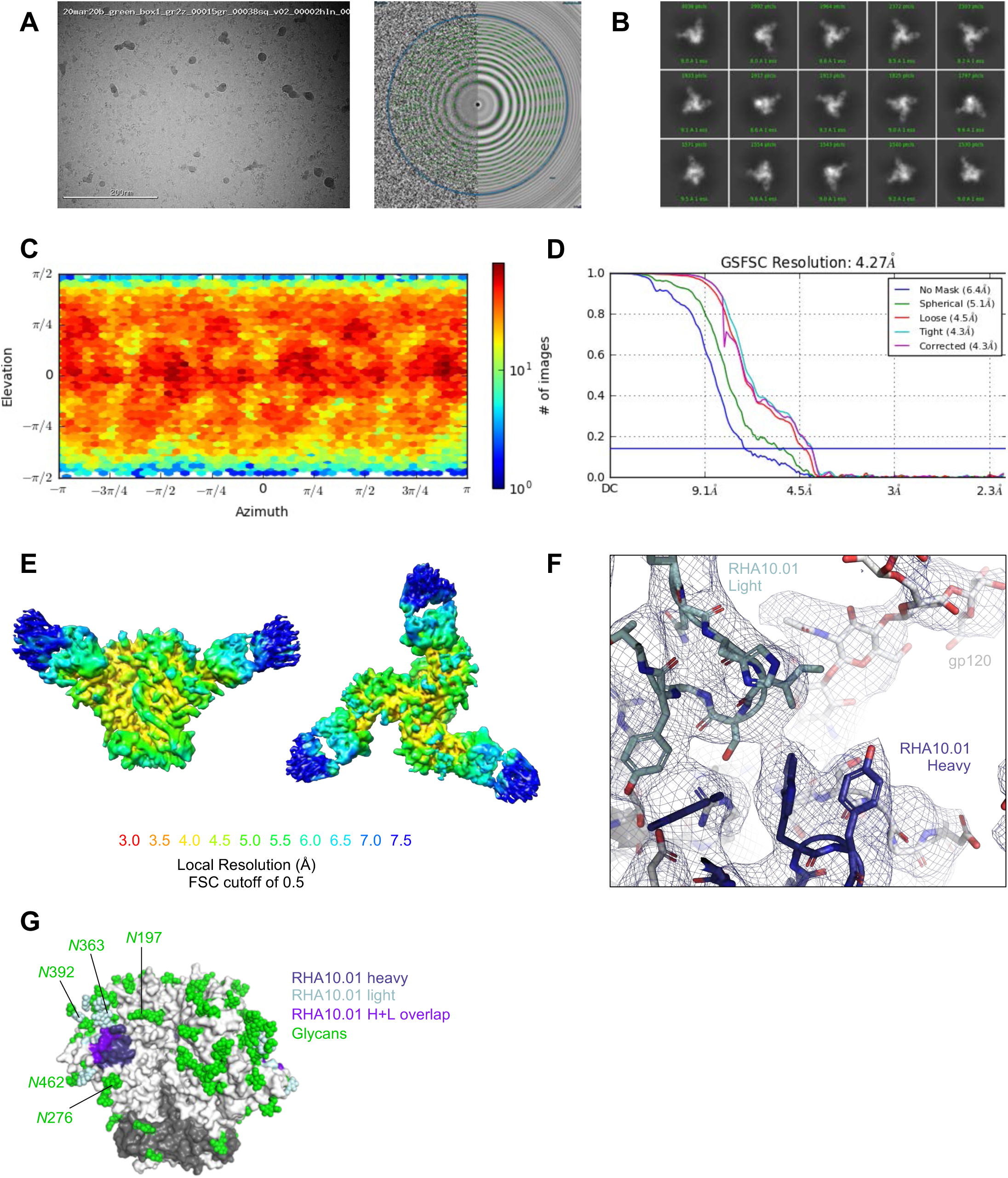
Cryo-EM Details of RHA10.01incomplex with HIV-1 Env BG505 DS-SOSIP. (A) Representative micrograph and CTF of the micrograph are shown. (B) Representative 2D class averages are shown. (C) The orientations of all particles used in the final refinement are shown as a heatmap. (D) The gold-standard fourier shell correlation resulted in a resolution of 4.27 Å using non-uniform refinement with C3 symmetry. (E) Local resolution of the full map is shown, generated with cryoSPARC using an FSC cutoff of 0.5. Two orientations are shown. (F) Example density is shown with the heavy and light chain highlighted. (G) Glycans resolved in cryo-EM model in relation to RHA10.01 footprint. Glycans in light blue are contacted by RHA10.01 light chain. N276 glycan is not contacted by RHA10.01.

**Figure S4.**
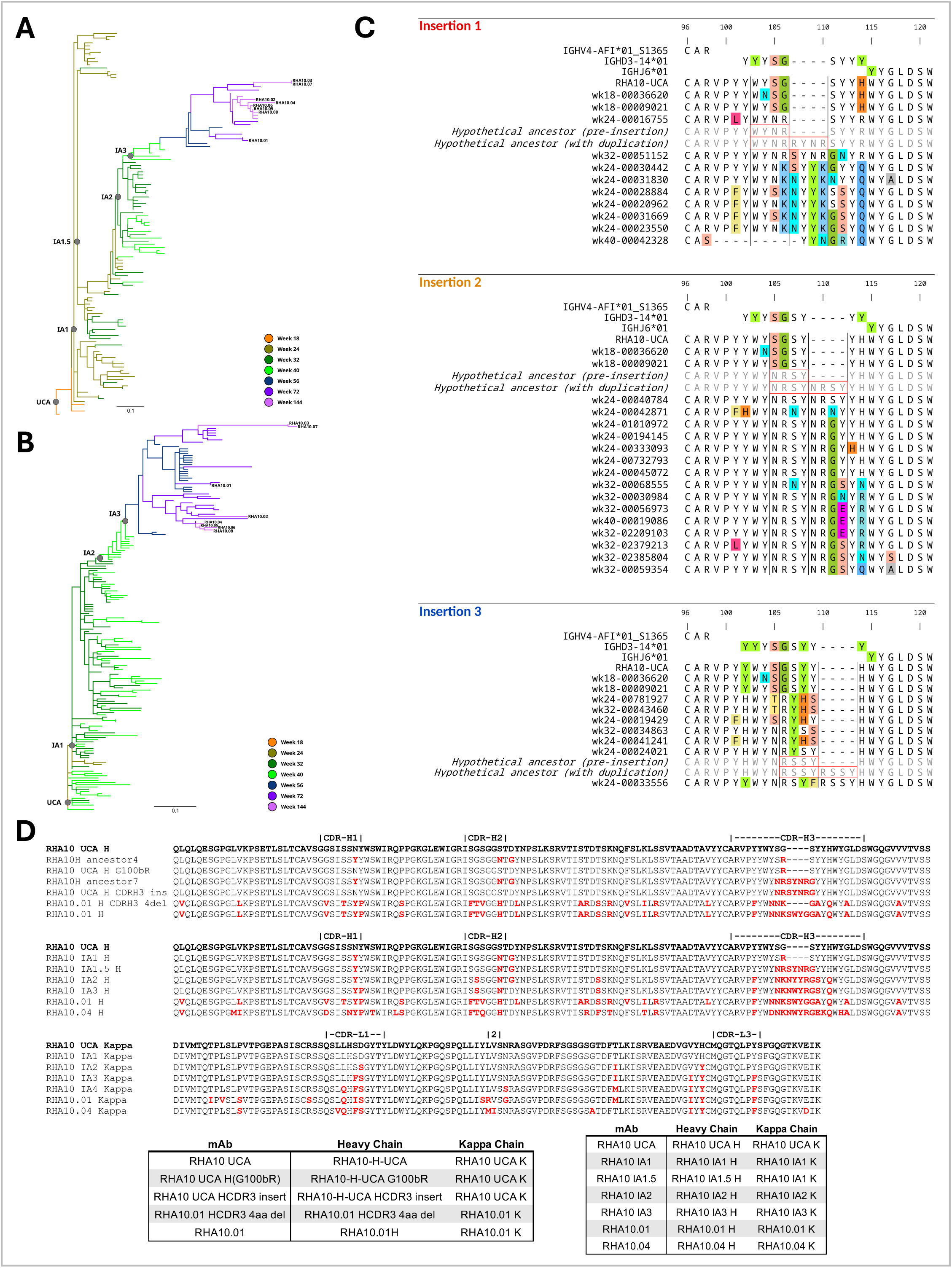
Lineage Tracing of the RHA10 lineage reveals a 4 amino-acid insertion in the CDR-H3. **A.**Phylogenetic tree of longitudinal RHA10 heavy chains from bulk-sorted and single-sorted IgG+ memory B cells colored by time point. **B.** Phylogenetic tree of longitudinal RHA10 kappa chains from bulk-sorted and single­sorted IgG+ memory B cells colored by time point. **C.** Amino acid alignment showing heavy chain junctions for possible independent insertion events. Each alignment shows germline V, D, and J segments, the RHA10 UCA heavy chain sequence, a selection of relevant members identified from bulk NGS, and a pair of hypothetical intermediate ancestors (gray text) before and after a 4 amino acid duplication (red outlines) producing each insertion. Disagreements with each post-insertion hypothetical ancestor are highlighted for each alignment. Given the similarity between adjacent sequence motifs post-insertion, the rapid lineage development between available sampled time points, and the inherent limited certainty of the phylogenetic inference, the true phylogeny may have included other insertions than the possibilities here. **D.** Amino acid alignments highlighting recombinant RHA10 insertion mutants. RHA10 ancestor4 and ancestor7 are inferred intermediates from the phylogenetic tree that we used to determine that G100bR occurred before the insertion and a less mutated version of the insertion, respectively. RHA10.01 contains the insertion but with additional SMH. Additionally, RHA10.01 CDR-H3 4aa del removes the 4 aa insertion but maintains the rest of the accumulated SHM outside of it. Heavy and kappa chain pairings for mAbs in Fig 3C are shown. Disagreements with the RHA10 UCA heavy and kappa chains are highlighted. RHA10 CDR-L2 is 3 amino acids long and is labeled as “2” in the alignment. GenBank accession numbers for the RHA10 UCA and inferred intermediate heavy and kappa chains are: PQ550586 – PQ550591, PQ550600 – PQ550605.

**Figure S5.**
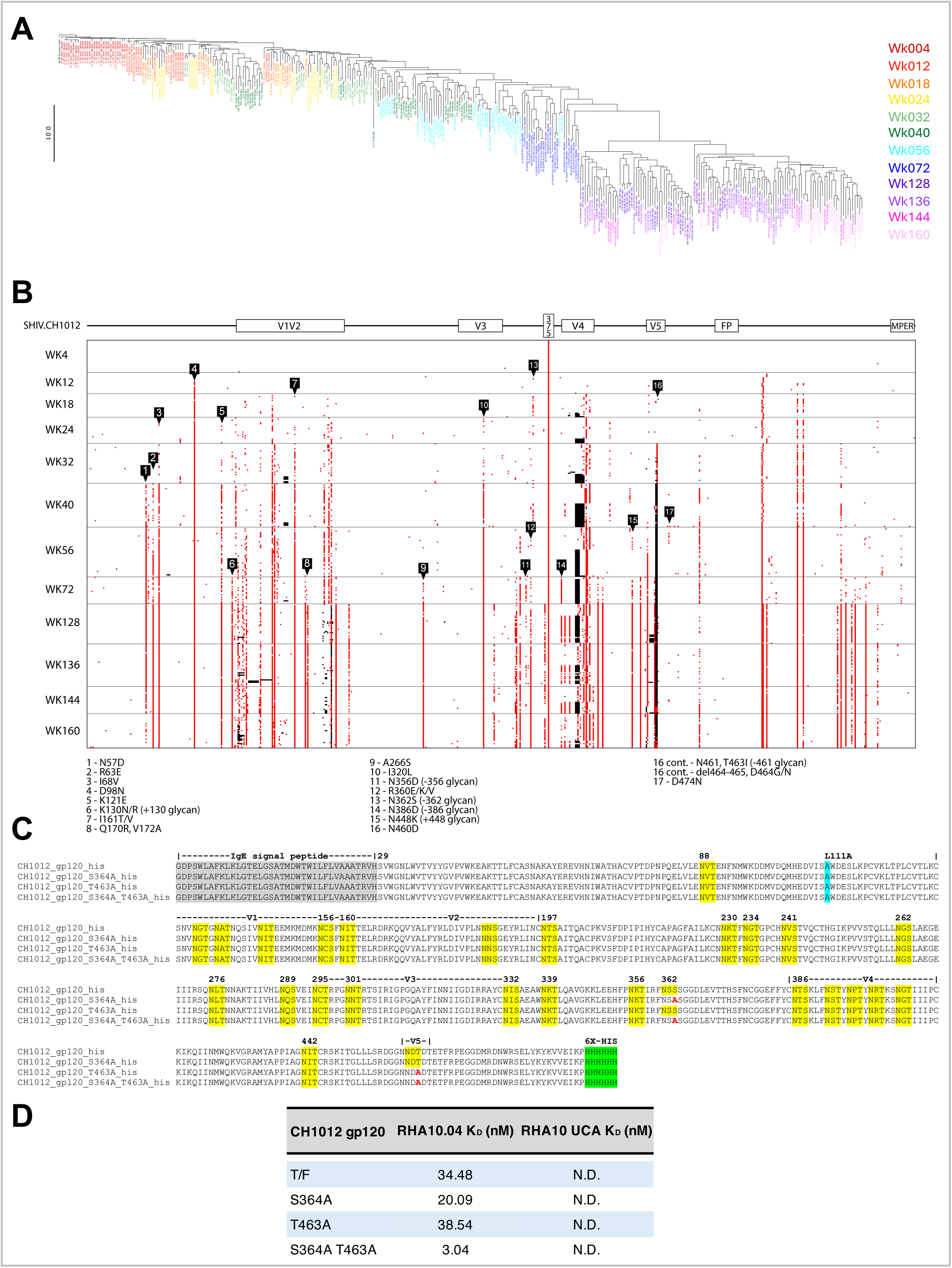
Single genome sequencing in macaque T681 reveals deletion of the N362 and N461 glycans prior to RHA10 elicitation. **A.** Phylogenetic tree of longitudinal viral Env nucleotide sequences from T681, colored by time point, rooted on the CH1012 T/F env. GenBank accession numbers: PQ574114 – PQ574448. **B.** Single viral genomes (rows) from RM T681 longitudinal samples aligned to the SHIV.CH1012 Env ectodomain sequence in a *Pixel* plot. Red, acquired mutations, black, insertion/deletions. **C.** Amino acid alignment of gp120 constructs used in this paper. PNGS highlighted in yellow, Env mutations away from T/F represented in bold red, residues identical to the T/F env represented in dashes. IgE signal peptide added to the N-terminus, L111A added to prevent oligomerization (Hoffenberg et al., 2013), and His-tag added to C-terminus. Genbank accession numbers: PP171710– PP171713 **D.** Binding K_D_s were determined using an Octet R8 protein analysis system (Sartorius). RHA10.04 or RHA10 UCA IgG was diluted in PBST with 1% BSA to a concentration of 5µg/ml and captured on anti-human IgG-F_c_ biosensors (Sartorius) for 300s. Association was measured by incubating the loaded sensors with gp120s for 300s, and dissociation was measured by incubating the sensors in PBST with 1% BSA for 600s. For gp120s, a starting concentration of 2µM followed by eight 3­fold dilutions was used for RHA10.04 affinity measurements, and a starting concentration of 50µM followed by four 2­fold dilutions was used for RHA10 UCA affinity measurements. Data was analyzed using Octet Analysis Studio Software (Sartorius).

**Figure S6.**
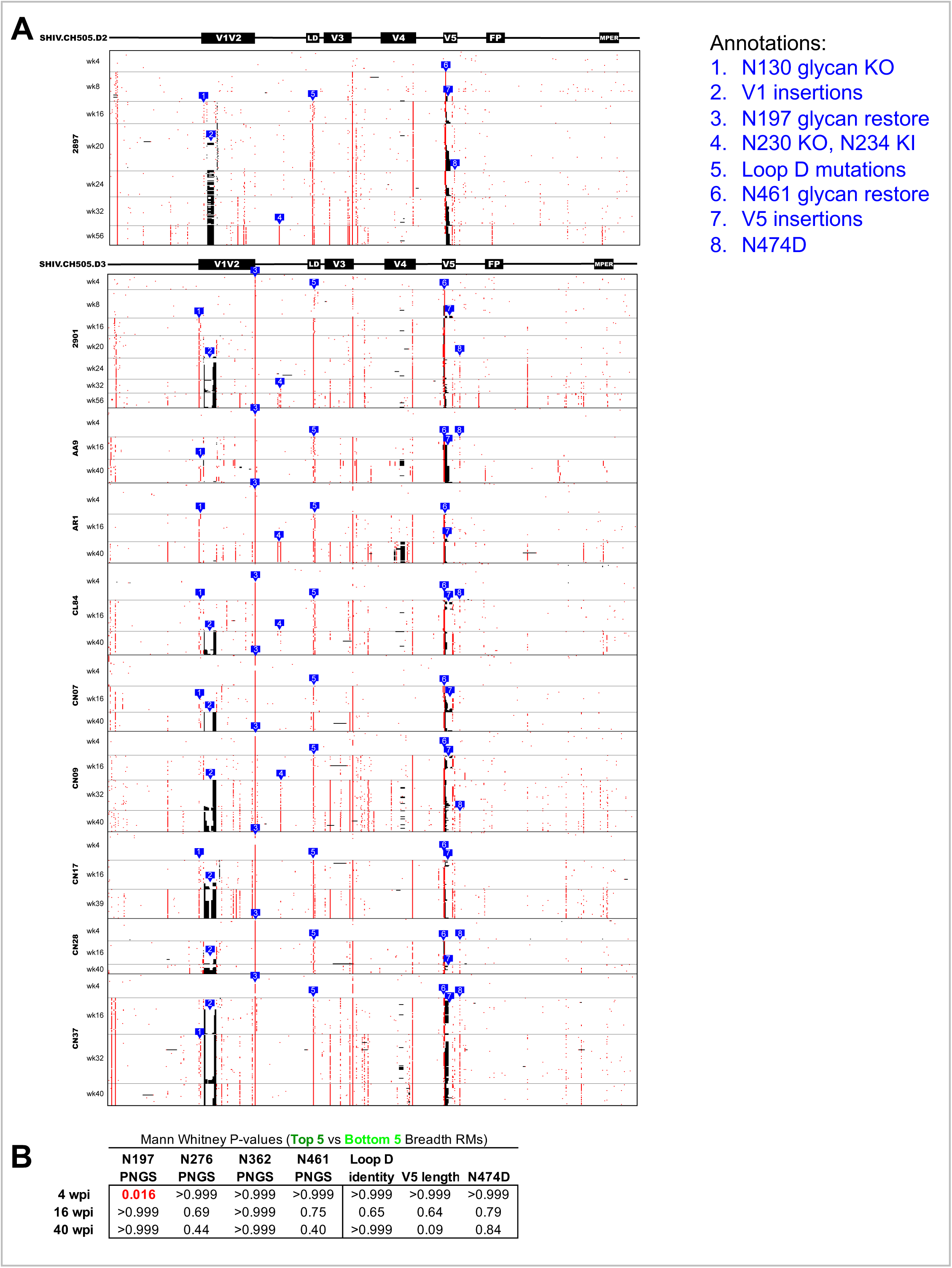
Single genome sequencing of SHIV.CH505.D2 and D3 infected macaques exhibits sequential glycan restoration and CD4bs epitope escape. **A.** Single viral genomes (rows) from SHIV.CH505.D2 and SHIV.CH505.D3 longitudinal samples aligned to the respective Env ectodomain sequence in a *Pixel* plot. Red, acquired mutations, black, insertion/deletions. GenBank accession numbers: PQ574449 – PQ575767. **B.** Summary of Mann-Whitney p-values comparing viral features at each time point between top 5 and bottom 5 SHIV.CH505.CD4bs.GH infected macaques

**Table S1.** RHA10.01 and RHA10.04 neutralization of a heterologous 119-virus panel. RHA10.01 and RHA10.04 were screened for neutralization on a 119-strain multiclade panel and their neutralizing IC50s in μg/mL are shown and colored by potency. Viruses organized by clade.

|  |  | Neut. IC50 (µg/ml) |  |
| --- | --- | --- | --- |
|  |  | RHA10.01 | RHA10.04 |
| Virus ID | Clade* | IC50 | IC50 |
| MS208.A1 | A | >50 | 37.405 |
| Q23.17 | A | >50 | >50 |
| Q461.e2 | A | >50 | >50 |
| Q769.d22 | A | >50 | 0.114 |
| Q259.d2.17 | A | 6.106 | 0.803 |
| Q842.d12 | A | 0.215 | 0.015 |
| 0260.v5.c36 | A | >50 | 6.593 |
| 3415.v1.c1 | A | >50 | >50 |
| 3365.v2.c20 | A | 1.092 | 0.108 |
| 191955_A11 | A (T/F) | >50 | >50 |
| 191084 B7-19 | A (T/F) | >50 | 0.030 |
| 9004SS_A3_4 | A (T/F) | >50 | >50 |
| 3301.v1.c24 | AC | >50 | 0.396 |
| 6041.v3.c23 | AC | >50 | >50 |
| 6540.v4.c1 | AC | 1.457 | >50 |
| 6545.v4.c1 | AC | >50 | >50 |
| 0815.v3.c3 | ACD | 4.595 | 0.013 |
| 3103.v3.c10 | ACD | >50 | >50 |
| 620345.c01 | CRF01_AE | >50 | >50 |
| CNE8 | CRF01_AE | >50 | 0.346 |
| C1080.c03 | CRF01_AE | >50 | >50 |
| R2184.c04 | CRF01_AE | >50 | 2.467 |
| R1166.c01 | CRF01_AE | >50 | 0.527 |
| R3265.c06 | CRF01_AE | >50 | >50 |
| C2101.c01 | CRF01_AE | >50 | 9.826 |
| C3347.c11 | CRF01_AE | 19.924 | 0.079 |
| C4118.c09 | CRF01_AE | >50 | 7.658 |
| CNE5 | CRF01_AE | 0.148 | 0.054 |
| BJOX009000.02.4 | CRF01_AE | >50 | >50 |
| BJOX015000.11.5 | CRF01_AE (T/F) | >50 | 3.296 |
| BJOX010000.06.2 | CRF01_AE (T/F) | 10.705 | 0.071 |
| BJOX025000.01.1 | CRF01_AE (T/F) | >50 | 0.025 |
| BJOX028000.10.3 | CRF01_AE (T/F) | >50 | >50 |
| T257-31 | CRF02_AG | >50 | >50 |
| 928-28 | CRF02_AG | >50 | >50 |
| 263-8 | CRF02_AG | >50 | 0.766 |
| T250-4 | CRF02_AG | >50 | >50 |
| T251-18 | CRF02_AG | >50 | >50 |
| T278-50 | CRF02_AG | >50 | >50 |
| T255-34 | CRF02_AG | >50 | >50 |
| 211-9 | CRF02_AG | >50 | >50 |
| 235-47 | CRF02_AG | >50 | >50 |
| 6535.3 | B | >50 | >50 |
| QH0692.42 | B | >50 | >50 |
| SC422661.8 | B | 0.916 | 0.216 |
| PVO.4 | B | >50 | 24.894 |
| TRO.11 | B | 5.421 | 1.391 |
| AC10.0.29 | B | 41.629 | >50 |
| RHPA4259.7 | B | 2.965 | 0.001 |
| THRO4156.18 | B | 3.479 | 0.267 |
| REJO4541.67 | B | >50 | >50 |
| TRJO4551.58 | B | >50 | 0.438 |
| WITO4160.33 | B | >50 | >50 |
| CAAN5342.A2 | B | >50 | >50 |
| WEAU_d15_410_787 | B (T/F) | >50 | 50.000 |
| 1006_11_C3_1601 | B (T/F) | >50 | >50 |
| 1054_07_TC4_1499 | B (T/F) | >50 | 2.354 |
| 1056_10_TA11_1826 | B (T/F) | 0.919 | 0.302 |
| 1012_11_TC21_3257 | B (T/F) | 0.175 | 0.082 |
| 6240_08_TA5_4622 | B (T/F) | >50 | 48.586 |
| 6244_13_B5_4576 | B (T/F) | >50 | >50 |
| 62357_14_D3_4589 | B (T/F) | >50 | 1.382 |
| SC05_8C11_2344 | B (T/F) | 2.042 | 0.541 |
|  |  | RHA10.01 | RHA10.04 |
| Virus ID | Clade* | IC50 | IC50 |
| CNE19 | BC | >50 | 0.022 |
| CNE20 | BC | 0.034 | 0.101 |
| CNE21 | BC | 24.748 | 1.284 |
| CNE17 | BC | >50 | >50 |
| CNE30 | BC | >50 | >50 |
| CNE52 | BC | >50 | >50 |
| CNE53 | BC | >50 | 1.632 |
| CNE58 | BC | >50 | 8.436 |
| Du156.12 | C | >50 | >50 |
| Du172.17 | C | >50 | >50 |
| Du422.1 | C | >50 | >50 |
| ZM197M.PB7 | C | 0.759 | 0.043 |
| ZM214M.PL15 | C | 0.07 | 0.058 |
| ZM233M.PB6 | C | >50 | >50 |
| ZM249M.PL1 | C | 26.929 | >50 |
| ZM53M.PB12 | C | >50 | 0.202 |
| ZM109F.PB4 | C | 0.121 | 1.202 |
| ZM135M.PL10a | C | >50 | >50 |
| CAP45.2.00.G3 | C | >50 | >50 |
| CAP210.2.00.E8 | C | >50 | >50 |
| HIV-001428-2.42 | C | >50 | 0.064 |
| HIV-0013095-2.11 | C | >50 | 0.195 |
| HIV-16055-2.3 | C | >50 | >50 |
| HIV-16845-2.22 | C | >50 | >50 |
| Ce1086_B2 | C (T/F) | >50 | 11.340 |
| Ce0393_C3 | C (T/F) | 50 | 13.454 |
| Ce1176_A3 | C (T/F) | >50 | 3.730 |
| Ce2010_F5 | C (T/F) | 10.73 | 0.405 |
| Ce0682_E4 | C (T/F) | >50 | >50 |
| Ce1172_H1 | C (T/F) | >50 | >50 |
| Ce2060_G9 | C (T/F) | 0.007 | 0.007 |
| Ce703010054_2A2 | C (T/F) | >50 | 1.423 |
| BF1266.431a | C (T/F) | >50 | >50 |
| 246F_C1G | C (T/F) | >50 | 2.312 |
| 249M_B10 | C (T/F) | 0.062 | 0.032 |
| ZM247v1(Rev-) | C (T/F) | >50 | >50 |
| 7030102001E5(Rev-) | C (T/F) | 9.724 | 0.197 |
| 1394C9G1(Rev-) | C (T/F) | >50 | 47.260 |
| Ce704809221_1B3 | C (T/F) | >50 | >50 |
| 3817.v2.c59 | CD | >50 | >50 |
| 6480.v4.c25 | CD | >50 | >50 |
| 6952.v1.c20 | CD | >50 | >50 |
| 6811.v7.c18 | CD | >50 | >50 |
| 89-F1_2_25 | CD | >50 | >50 |
| 3016.v5.c45 | D | 0.424 | 0.030 |
| A07412M1.vrc12 | D | 41.821 | 0.096 |
| 231965.c01 | D | >50 | >50 |
| 231966.c02 | D | 0.218 | 0.016 |
| 6405.v4.c34 | D | >50 | >50 |
| X1193_c1 | G | >50 | 0.434 |
| P0402_c2_11 | G | >50 | >50 |
| X1254_c3 | G | >50 | >50 |
| X2088_c9 | G | >50 | >50 |
| X2131_C1_B5 | G | 45.962 | 0.151 |
| P1981_C5_3 | G | >50 | 42.201 |
| X1632_S2_B10 | G | 0.1 | 0.011 |
| MuLV | Neg. Control | >50 | >50 |

\* (T/F): Transmitted / Founder Virus

**Table S2.** Cryo-EM Data Collection Statistics for RHA10.01 bound to BG505 DS-SOSIP.664.

|  | RHA10.01 Fab with<br>BG505 DS-SOSIP.664 |
| --- | --- |
| EMDB ID | 47000 |
| PDB ID | 9DMB |
| <u>Data collection</u> |  |
| Microscope | FEI Titan Krios |
| Voltage (kV) | 300 |
| Electron dose (e<sup>-</sup>/Å<sup>2</sup>) | 70.73 |
| Detector | Gatan K3 |
| Pixel Size (Å) | 1.083 |
| Defocus Range (µm) | -0.8 to -2.5 |
| Magnification | 81,000 |
| <u>Reconstruction</u> |  |
| Software | CryoSparc v3 |
| Particles | 81,730 |
| Symmetry | C3 |
| Box size (pix) | 420 |
| Resolution (Å) (FSC<sub>0.143</sub>) | 4.27 |
| <u>Refinement</u> |  |
| Software | Phenix v1.21 |
| Protein residues | 2448 |
| Chimera CC | 0.75 |
| EMRinger Score (resolve map) | 1.36 |
| R.m.s. deviations |  |
| Bond lengths (Å) | 0.003 |
| Bond angles (°) | 0.569 |
| <u>Validation</u> |  |
| Molprobity score | 1.66 |
| Clash score | 4.71 |
| Favored rotamers (%) | 99.28 |
| Ramachandran |  |
| Favored regions (%) | 93.62 |
| Disallowed regions (%) | 0.25 |

**Supplemental Table 4.** T681 viral NGS read processing breakdown and PNGS frequencies. A. NGS read processing breakdown across sample time points B. Raw counts and percentages of NGS processed reads that lack either or both 362 and 461 PNGS. Viral NGS from macaque T681 was deposited to NCBI SRA under PRJNA1180587.

| A | WK4 (D28) |  |  | WK8 (D56) |  |  | WK12 (D84) |  |  | WK18 (D126) |  |  | WK24 (D166) |  |  |
| --- | --- | --- | --- | --- | --- | --- | --- | --- | --- | --- | --- | --- | --- | --- | --- |
|  | Number | % of parent | % of total | Number | % of parent | % of total | Number | % of parent | % of total | Number | % of parent | % of total | Number | % of parent | % of total |
| Paired Reads | 244,869 | n.a. | 100.0 | 147,530 | n.a. | 100.0 | 364,987 | n.a. | 100.0 | 202,590 | n.a. | 100.0 | 238,275 | n.a. | 100.0 |
| Merged Reads | 190,084 | 77.6 | 77.6 | 110,117 | 74.6 | 74.6 | 288,569 | 79.1 | 79.1 | 169,669 | 83.7 | 83.7 | 184,956 | 77.6 | 77.6 |
| Deduplicated Reads | 126,429 | 66.5 | 51.6 | 75,637 | 68.7 | 51.3 | 193,998 | 67.2 | 53.2 | 134,412 | 79.2 | 66.3 | 148,819 | 80.5 | 62.5 |
| Reads assembled to CH1012 Env | 124,656 | 98.6 | 50.9 | 75,492 | 99.8 | 51.2 | 171,362 | 88.3 | 47.0 | 100,221 | 74.6 | 49.5 | 130,649 | 87.8 | 54.8 |
| Reads that span both glycans | 121,563 | 97.5 | 49.6 | 74,848 | 99.1 | 50.7 | 162,339 | 94.7 | 44.5 | 96,418 | 96.2 | 47.6 | 126,970 | 97.2 | 53.3 |
| Length filtered translations | 121,183 | 99.7 | 49.5 | 74,848 | 100.0 | 50.7 | 161,510 | 99.5 | 44.3 | 96,406 | 100.0 | 47.6 | 126,199 | 99.4 | 53.0 |

| B | WK4 (D28) |  | WK8 (D56) |  | WK12 (D84) |  | WK18 (D126) |  | WK24 (D166) |  |
| --- | --- | --- | --- | --- | --- | --- | --- | --- | --- | --- |
|  | Number | % | Number | % | Number | % | Number | % | Number | % |
| Both glycans intact (T/F) | 116,928 | 96.49 | 71,173 | 95.09 | 138,207 | 85.57 | 46,560 | 48.30 | 89,345 | 70.80 |
| N362 glycan knockout | 1,890 | 1.56 | 1,722 | 2.30 | 14,862 | 9.20 | 33,713 | 34.97 | 29,596 | 23.45 |
| N461 glycan knockout | 2,295 | 1.89 | 1,822 | 2.43 | 7,731 | 4.79 | 10,609 | 11.00 | 5,901 | 4.68 |
| Dual glycan knockout | 69 | 0.06 | 131 | 0.18 | 710 | 0.44 | 5,524 | 5.73 | 1,357 | 1.08 |
| Total Sequences | 121,182 | 100.0 | 74,848 | 100.0 | 161,510 | 100.0 | 96,406 | 100.0 | 126,199 | 100.0 |

## REFERENCES

1. Haynes, B.F., Burton, D.R., and Mascola, J.R. (2019). Multiple roles for HIV broadly neutralizing antibodies. Sci. Transl. Med. 11, eaaz2686. 10.1126/scitranslmed.aaz2686.

2. Keele, B.F., Giorgi, E.E., Salazar-Gonzalez, J.F., Decker, J.M., Pham, K.T., Salazar, M.G., Sun, C., Grayson, T., Wang, S., Li, H., et al. (2008). Identification and characterization of transmitted and early founder virus envelopes in primary HIV-1 infection. Proc. Natl. Acad. Sci. 10.1073/pnas.0802203105.

3. Li, H., Wang, S., Kong, R., Ding, W., Lee, F.-H., Parker, Z., Kim, E., Learn, G.H., Hahn, P., Policicchio, B., et al. (2016). Envelope residue 375 substitutions in simian–human immunodeficiency viruses enhance CD4 binding and replication in rhesus macaques. Proc. Natl. Acad. Sci. 113, E3413–E3422. 10.1073/pnas.1606636113.

4. Roark, R.S., Li, H., Williams, W.B., Chug, H., Mason, R., Gorman, J., Wang, S., Lee, F., Rando, J., Bonsignori, M., et al. (2020). Recapitulation of HIV-1 Env-Antibody Coevolution in Macaques Leading to Neutralization Breadth. Science 2638, 1–32.

5. Zhou, T., Lynch, R.M., Chen, L., Acharya, P., Wu, X., Doria-Rose, N.A., Joyce, M.G., Lingwood, D., Soto, C., Bailer, R.T., et al. (2015). Structural Repertoire of HIV-1-Neutralizing Antibodies Targeting the CD4 Supersite in 14 Donors. Cell 161, 1280–1292.

6. Huang, J., Kang, B.H., Ishida, E., Zhou, T., Griesman, T., Sheng, Z., Wu, F., Doria-Rose, N.A., Zhang, B., McKee, K., et al. (2016). Identification of a CD4-Binding-Site Antibody to HIV that Evolved Near-Pan Neutralization Breadth. Immunity 45, 1108–1121. 10.1016/j.immuni.2016.10.027.

7. Liao, H.X., Lynch, R., Zhou, T., Gao, F., Munir Alam, S., Boyd, S.D., Fire, A.Z., Roskin, K.M., Schramm, C.A., Zhang, Z., et al. (2013). Co-evolution of a broadly neutralizing HIV-1 antibody and founder virus. Nature 496, 469–476. 10.1038/nature12053.

8. Caniels, T.G., Medina-Ramìrez, M., Zhang, S., Kratochvil, S., Xian, Y., Koo, J.-H., Derking, R., Samsel, J., Schooten, J. van, Pecetta, S., et al. (2024). Germline-targeting HIV vaccination induces neutralizing antibodies to the CD4 binding site. Sci. Immunol. 9, eadk9550. 10.1126/sciimmunol.adk9550.

9. Nelson, A.N., Shen, X., Vekatayogi, S., Zhang, S., Ozorowski, G., Dennis, M., Sewall, L.M., Milligan, E., Davis, D., Cross, K.A., et al. (2024). Immunization with germ line–targeting SOSIP trimers elicits broadly neutralizing antibody precursors in infant macaques. Sci. Immunol. 9, eadm7097. 10.1126/sciimmunol.adm7097.

10. Jardine, J., Julien, J.-P., Menis, S., Ota, T., Kalyuzhniy, O., McGuire, A., Sok, D., Huang, P.-S., MacPherson, S., Jones, M., et al. (2013). Rational HIV immunogen design to target specific germline B cell receptors. Science 340, 711–716. 10.1126/science.1234150.

11. Jardine, J.G., Kulp, D.W., Havenar-Daughton, C., Sarkar, A., Briney, B., Sok, D., Sesterhenn, F., Ereño-Orbea, J., Kalyuzhniy, O., Deresa, I., et al. (2016). HIV-1 broadly neutralizing antibody precursor B cells revealed by germline-targeting immunogen. Science 351, 1458–1463. 10.1126/science.aad9195.

12. Leggat, D.J., Cohen, K.W., Willis, J.R., Fulp, W.J., deCamp, A.C., Kalyuzhniy, O., Cottrell, C.A., Menis, S., Finak, G., Ballweber-Fleming, L., et al. Vaccination induces HIV broadly neutralizing antibody precursors in humans. Science 378, eadd6502. 10.1126/science.add6502.

13. Vázquez Bernat, N., Corcoran, M., Nowak, I., Kaduk, M., Castro Dopico, X., Narang, S., Maisonasse, P., Dereuddre-Bosquet, N., Murrell, B., and Karlsson Hedestam, G.B. (2021). Rhesus and cynomolgus macaque immunoglobulin heavy-chain genotyping yields comprehensive databases of germline VDJ alleles. Immunity 54, 355–366.e4. 10.1016/j.immuni.2020.12.018.

14. Giudicelli, V., Duroux, P., Ginestoux, C., Folch, G., Jabado-Michaloud, J., Chaume, D., and Lefranc, M.-P. (2006). IMGT/LIGM-DB, the IMGT® comprehensive database of immunoglobulin and T cell receptor nucleotide sequences. Nucleic Acids Res. 34, D781–D784. 10.1093/nar/gkj088.

15. Saunders, K.O., Counts, J., Thakur, B., Stalls, V., Edwards, R., Manne, K., Lu, X., Mansouri, K., Chen, Y., Parks, R., et al. (2024). Vaccine induction of CD4-mimicking HIV-1 broadly neutralizing antibody precursors in macaques. Cell 187, 79–94.e24. 10.1016/j.cell.2023.12.002.

16. Landais, E., and Moore, P.L. (2018). Development of broadly neutralizing antibodies in HIV-1 infected elite neutralizers. Retrovirology 15, 1–14. 10.1186/s12977-018-0443-0.

17. Zhou, T., Doria-Rose, N.A., Cheng, C., Stewart-Jones, G.B.E., Chuang, G.Y., Chambers, M., Druz, A., Geng, H., McKee, K., Kwon, Y.D., et al. (2017). Quantification of the Impact of the HIV-1-Glycan Shield on Antibody Elicitation. Cell Rep. 19, 719–732. 10.1016/j.celrep.2017.04.013.

18. Li, H., Wang, S., Lee, F.-H., Roark, R.S., Murphy, A.I., Smith, J., Zhao, C., Rando, J., Chohan, N., Ding, Y., et al. (2021). New SHIVs and Improved Design Strategy for Modeling HIV-1 Transmission, Immunopathogenesis, Prevention and Cure. J. Virol. 95, JVI.00071-21. 10.1128/JVI.00071-21.

19. Wu, X., Yang, Z.-Y., Li, Y., Hogerkorp, C.-M., Schief, W.R., Seaman, M.S., Zhou, T., Schmidt, S.D., Wu, L., Xu, L., et al. (2010). Rational Design of Envelope Identifies Broadly Neutralizing Human Monoclonal Antibodies to HIV-1. Science 329, 856–861. 10.1126/science.1187659.

20. Kwon, Y.D., Pancera, M., Acharya, P., Georgiev, I.S., Crooks, E.T., Gorman, J., Joyce, M.G., Guttman, M., Ma, X., Narpala, S., et al. (2015). Crystal structure, conformational fixation and entry-related interactions of mature ligand-free HIV-1 Env. Nat. Struct. Mol. Biol. 22, 522–531. 10.1038/nsmb.3051.

21. Mason, R.D., Welles, H.C., Adams, C., Chakrabarti, B.K., Gorman, J., Zhou, T., Nguyen, R., O’Dell, S., Lusvarghi, S., Bewley, C.A., et al. (2016). Targeted Isolation of Antibodies Directed against Major Sites of SIV Env Vulnerability. PLoS Pathog. 12, e1005537. 10.1371/journal.ppat.1005537.

22. Yoon, H., Macke, J., West, A.P.J., Foley, B., Bjorkman, P.J., Korber, B., and Yusim, K. (2015). CATNAP: a tool to compile, analyze and tally neutralizing antibody panels. Nucleic Acids Res. 43, W213–219. 10.1093/nar/gkv404.

23. Corey, L., Gilbert, P.B., Juraska, M., Montefiori, D.C., Morris, L., Karuna, S.T., Edupuganti, S., Mgodi, N.M., deCamp, A.C., Rudnicki, E., et al. (2021). Two Randomized Trials of Neutralizing Antibodies to Prevent HIV-1 Acquisition. N. Engl. J. Med. 384, 1003–1014. 10.1056/NEJMoa2031738.

24. Pauthner, M.G., Nkolola, J.P., Havenar-Daughton, C., Murrell, B., Reiss, S.M., Bastidas, R., Prévost, J., Nedellec, R., von Bredow, B., Abbink, P., et al. (2019). Vaccine-Induced Protection from Homologous Tier 2 SHIV Challenge in Nonhuman Primates Depends on Serum-Neutralizing Antibody Titers. Immunity 50, 241–252.e6. 10.1016/j.immuni.2018.11.011.

25. Saunders, K.O., Edwards, R.J., Tilahun, K., Manne, K., Lu, X., Cain, D.W., Wiehe, K., Williams, W.B., Mansouri, K., Hernandez, G.E., et al. Stabilized HIV-1 envelope immunization induces neutralizing antibodies to the CD4bs and protects macaques against mucosal infection. Sci. Transl. Med. 14, eabo5598. 10.1126/scitranslmed.abo5598.

26. Corti, D., Langedijk, J.P.M., Hinz, A., Seaman, M.S., Vanzetta, F., Fernandez-Rodriguez, B.M., Silacci, C., Pinna, D., Jarrossay, D., Balla-Jhagjhoorsingh, S., et al. (2010). Analysis of Memory B Cell Responses and Isolation of Novel Monoclonal Antibodies with Neutralizing Breadth from HIV-1-Infected Individuals. PLOS ONE 5, 1–15. 10.1371/journal.pone.0008805.

27. Freund, N.T., Horwitz, J.A., Nogueira, L., Sievers, S.A., Scharf, L., Scheid, J.F., Gazumyan, A., Liu, C., Velinzon, K., Goldenthal, A., et al. (2015). A New Glycan-Dependent CD4-Binding Site Neutralizing Antibody Exerts Pressure on HIV-1 In Vivo. PLOS Pathog. 11, 1–19. 10.1371/journal.ppat.1005238.

28. Cottrell, C.A., Manne, K., Kong, R., Wang, S., Zhou, T., Chuang, G.Y., Edwards, R.J., Henderson, R., Janowska, K., Kopp, M., et al. (2021). Structural basis of glycan276-dependent recognition by HIV-1 broadly neutralizing antibodies. Cell Rep. 37, 109922. 10.1016/j.celrep.2021.109922.

29. Corcoran, M.M., Phad, G.E., Bernat, N.V., Stahl-Hennig, C., Sumida, N., Persson, M.A.A., Martin, M., and Hedestam, G.B.K. (2016). Production of individualized v gene databases reveals high levels of immunoglobulin genetic diversity. Nat. Commun. 7. 10.1038/ncomms13642.

30. Schramm, C.A., Sheng, Z., Zhang, Z., Mascola, J.R., Kwong, P.D., and Shapiro, L. (2016). SONAR: A high-throughput pipeline for inferring antibody ontogenies from longitudinal sequencing of B cell transcripts. Front. Immunol. 7, 1–10. 10.3389/fimmu.2016.00372.

31. Rudicell, R.S., Kwon, Y.D., Ko, S.-Y., Pegu, A., Louder, M.K., Georgiev, I.S., Wu, X., Zhu, J., Boyington, J.C., Chen, X., et al. (2014). Enhanced potency of a broadly neutralizing HIV-1 antibody in vitro improves protection against lentiviral infection in vivo. J. Virol. 88, 12669–12682. 10.1128/JVI.02213-14.

32. Dubrovskaya, V., Guenaga, J., de Val, N., Wilson, R., Feng, Y., Movsesyan, A., Karlsson Hedestam, G.B., Ward, A.B., and Wyatt, R.T. (2017). Targeted N-glycan deletion at the receptor-binding site retains HIV Env NFL trimer integrity and accelerates the elicited antibody response. PLoS Pathog. 13, 1–30. 10.1371/journal.ppat.1006614.

33. McGuire, A.T., Hoot, S., Dreyer, A.M., Lippy, A., Stuart, A., Cohen, K.W., Jardine, J., Menis, S., Scheid, J.F., West, A.P., et al. (2013). Engineering HIV envelope protein to activate germline B cell receptors of broadly neutralizing anti-CD4 binding site antibodies. J. Exp. Med. 210, 655–663. 10.1084/jem.20122824.

34. Medina-Ramírez, M., Garces, F., Escolano, A., Skog, P., de Taeye, S.W., Del Moral-Sanchez, I., McGuire, A.T., Yasmeen, A., Behrens, A.-J., Ozorowski, G., et al. (2017). Design and crystal structure of a native-like HIV-1 envelope trimer that engages multiple broadly neutralizing antibody precursors in vivo. J. Exp. Med. 214, 2573–2590. 10.1084/jem.20161160.

35. Briney, B., Sok, D., Jardine, J.G., Kulp, D.W., Skog, P., Menis, S., Jacak, R., Kalyuzhniy, O., de Val, N., Sesterhenn, F., et al. (2016). Tailored Immunogens Direct Affinity Maturation toward HIV Neutralizing Antibodies. Cell 166, 1459–1470.e11. 10.1016/j.cell.2016.08.005.

36. Saunders, K.O., Wiehe, K., Tian, M., Acharya, P., Bradley, T., Alam, S.M., Go, E.P., Scearce, R., Sutherland, L., Henderson, R., et al. (2019). Targeted selection of HIV-specific antibody mutations by engineering B cell maturation. Science 366. 10.1126/science.aay7199.

37. Gristick, H.B., Hartweger, H., Loewe, M., van Schooten, J., Ramos, V., Oliveira, T.Y., Nishimura, Y., Koranda, N.S., Wall, A., Yao, K.-H., et al. (2023). CD4 binding site immunogens elicit heterologous anti-HIV-1 neutralizing antibodies in transgenic and wild-type animals. Sci. Immunol. 8, eade6364. 10.1126/sciimmunol.ade6364.

38. West, A.P.J., Diskin, R., Nussenzweig, M.C., and Bjorkman, P.J. (2012). Structural basis for germ-line gene usage of a potent class of antibodies targeting the CD4-binding site of HIV-1 gp120. Proc. Natl. Acad. Sci. U. S. A. 109, E2083–2090. 10.1073/pnas.1208984109.

39. Klasse, P.J., LaBranche, C.C., Ketas, T.J., Ozorowski, G., Cupo, A., Pugach, P., Ringe, R.P., Golabek, M., van Gils, M.J., Guttman, M., et al. (2016). Sequential and Simultaneous Immunization of Rabbits with HIV-1 Envelope Glycoprotein SOSIP.664 Trimers from Clades A, B and C. PLOS Pathog. 12, 1–31. 10.1371/journal.ppat.1005864.

40. McCoy, L.E., Gils, M.J. van, Ozorowski, G., Messmer, T., Briney, B., Voss, J.E., Kulp, D.W., Macauley, M.S., Sok, D., Pauthner, M., et al. (2016). Holes in the Glycan Shield of the Native HIV Envelope Are a Target of Trimer-Elicited Neutralizing Antibodies. Cell Rep. 16, 2327–2338. 10.1016/j.celrep.2016.07.074.

41. Klasse, P.J., Ketas, T.J., Cottrell, C.A., Ozorowski, G., Debnath, G., Camara, D., Francomano, E., Pugach, P., Ringe, R.P., LaBranche, C.C., et al. (2018). Epitopes for neutralizing antibodies induced by HIV-1 envelope glycoprotein BG505 SOSIP trimers in rabbits and macaques. PLoS Pathog. 14, 1–20. 10.1371/journal.ppat.1006913.

42. van Schooten, J., van Haaren, M.M., Li, H., McCoy, L.E., Havenar-Daughton, C., Cottrell, C.A., Burger, J.A., van der Woude, P., Helgers, L.C., Tomris, I., et al. (2021). Antibody responses induced by SHIV infection are more focused than those induced by soluble native HIV-1 envelope trimers in non-human primates. PLOS Pathog. 17, 1–22. 10.1371/journal.ppat.1009736.

43. Ringe, R.P., Pugach, P., Cottrell, C.A., LaBranche, C.C., Seabright, G.E., Ketas, T.J., Ozorowski, G., Kumar, S., Schorcht, A., Gils, M.J. van, et al. (2019). Closing and Opening Holes in the Glycan Shield of HIV-1 Envelope Glycoprotein SOSIP Trimers Can Redirect the Neutralizing Antibody Response to the Newly Unmasked Epitopes. J. Virol. 93, 10.1128/jvi.01656-18. 10.1128/jvi.01656-18.

44. Hatziioannou, T., Del Prete, G.Q., Keele, B.F., Estes, J.D., McNatt, M.W., Bitzegeio, J., Raymond, A., Rodriguez, A., Schmidt, F., Mac Trubey, C., et al. (2014). HIV-1–induced AIDS in monkeys. Science 344, 1401–1405. 10.1126/science.1250761.

45. Roark, R.S., Habib, R., Gorman, J., Li, H., Connell, A.J., Bonsignori, M., Guo, Y., Hogarty, M.P., Olia, A.S., Sowers, K., et al. (2024). HIV-1 neutralizing antibodies in SHIV-infected macaques recapitulate structurally divergent modes of human V2 apex recognition with a single D gene. bioRxiv, 2024.06.11.598384. 10.1101/2024.06.11.598384.

46. Li, Y., O’Dell, S., Walker, L.M., Wu, X., Guenaga, J., Feng, Y., Schmidt, S.D., McKee, K., Louder, M.K., Ledgerwood, J.E., et al. (2011). Mechanism of neutralization by the broadly neutralizing HIV-1 monoclonal antibody VRC01. J. Virol. 85, 8954–8967. 10.1128/JVI.00754-11.

47. Gristick, H.B., von Boehmer, L., West, A.P.J., Schamber, M., Gazumyan, A., Golijanin, J., Seaman, M.S., Fätkenheuer, G., Klein, F., Nussenzweig, M.C., et al. (2016). Natively glycosylated HIV-1 Env structure reveals new mode for antibody recognition of the CD4-binding site. Nat. Struct. Mol. Biol. 23, 906–915. 10.1038/nsmb.3291.

48. Bonsignori, M., Zhou, T., Sheng, Z., Chen, L., Gao, F., Joyce, M.G., Ozorowski, G., Chuang, G.Y., Schramm, C.A., Wiehe, K., et al. (2016). Maturation Pathway from Germline to Broad HIV-1 Neutralizer of a CD4-Mimic Antibody. Cell 165, 449–463. 10.1016/j.cell.2016.02.022.

49. Gao, F., Bonsignori, M., Liao, H.X., Kumar, A., Xia, S.M., Lu, X., Cai, F., Hwang, K.K., Song, H., Zhou, T., et al. (2014). Cooperation of B cell lineages in induction of HIV-1-broadly neutralizing antibodies. Cell 158, 481–491. 10.1016/j.cell.2014.06.022.

50. Wei, X., Decker, J.M., Wang, S., Hui, H., Kappes, J.C., Wu, X., Salazar-Gonzalez, J.F., Salazar, M.G., Kilby, J.M., Saag, M.S., et al. (2003). Antibody neutralization and escape by HIV-1. Nature 837, 307–312.

51. Bar, K.J., Tsao, C., Iyer, S.S., Decker, J.M., Yang, Y., Bonsignori, M., Chen, X., Hwang, K.- K., Montefiori, D.C., Liao, H.-X., et al. (2012). Early Low-Titer Neutralizing Antibodies Impede HIV-1 Replication and Select for Virus Escape. PLOS Pathog. 8, 1–20. 10.1371/journal.ppat.1002721.

52. Bonsignori, M., Meyerhoff, R.R., Bradley, T., Wiehe, K., Alam, S.M., Williams, W.B., Liao, H.-X., Gao, F., Haynes, B.F., Hwang, K.-K., et al. (2017). Staged induction of HIV-1 glycan-dependent broadly neutralizing antibodies. Sci. Transl. Med. 9. 10.1126/scitranslmed.aai7514.

53. Pegu, A., Lovelace, S.E., DeMouth, M.E., Cully, M.D., Morris, D.J., Li, Y., Wang, K., Schmidt, S.D., Choe, M., Liu, C., et al. (2024). Antibodies targeting the fusion peptide on the HIV envelope provide protection to rhesus macaques against mucosal SHIV challenge. Sci. Transl. Med. 16, eadh9039. 10.1126/scitranslmed.adh9039.

54. Hua Wang, Cheng Cheng, James L. Dal Santo, Chen-Hsiang Shen, Tatsiana Bylund, Amy R. Henry, Colin A. Howe, Juyun Hwang, Nicholas C. Morano, Daniel J. Morris, et al. (In Press). Potent and broad HIV-1 neutralization in fusion peptide-primed SHIV-infected macaques. Cell. 10.1016/j.cell.2024.10.003.

55. Wagh, K., Kreider, E.F., Li, Y., Barbian, H.J., Learn, G.H., Giorgi, E., Hraber, P.T., Decker, T.G., Smith, A.G., Gondim, M.V., et al. (2018). Completeness of HIV-1 Envelope Glycan Shield at Transmission Determines Neutralization Breadth. Cell Rep. 25, 893–908.e7. 10.1016/j.celrep.2018.09.087.

56. Sundling, C., Li, Y., Huynh, N., Poulsen, C., Wilson, R., O’Dell, S., Feng, Y., Mascola, J.R., Wyatt, R.T., and Karlsson Hedestam, G.B. (2012). High-resolution definition of vaccine-elicited B cell responses against the HIV primary receptor binding site. Sci. Transl. Med. 4, 142ra96. 10.1126/scitranslmed.3003752.

57. Habib, R., Solieva, S.O., Lin, Z.J., Ghosh, S., Bayruns, K., Singh, M., Agostino, C.J., Tursi, N.J., Sowers, K.J., Huang, J., et al. (2024). Deep Mining of the Human Antibody Repertoire Identifies Frequent and Immunogenetically Diverse CDRH3 Topologies Targetable by Vaccination. bioRxiv, 2024.10.04.616739. 10.1101/2024.10.04.616739.

58. Wang, Z., Barnes, C.O., Gautam, R., Cetrulo Lorenzi, J.C., Mayer, C.T., Oliveira, T.Y., Ramos, V., Cipolla, M., Gordon, K.M., Gristick, H.B., et al. (2020). A broadly neutralizing macaque monoclonal antibody against the HIV-1 V3-Glycan patch. eLife 9, e61991. 10.7554/eLife.61991.

59. Steichen, J.M., Phung, I., Salcedo, E., Ozorowski, G., Willis, J.R., Baboo, S., Liguori, A., Cottrell, C.A., Torres, J.L., Madden, P.J., et al. Vaccine priming of rare HIV broadly neutralizing antibody precursors in nonhuman primates. Science 384, eadj8321. 10.1126/science.adj8321.

60. Escolano, A., Gristick, H.B., Gautam, R., DeLaitsch, A.T., Abernathy, M.E., Yang, Z., Wang, H., Hoffmann, M.A.G., Nishimura, Y., Wang, Z., et al. (2021). Sequential immunization of macaques elicits heterologous neutralizing antibodies targeting the V3-glycan patch of HIV-1 Env. Sci. Transl. Med. 13, eabk1533. 10.1126/scitranslmed.abk1533.

61. Zhang, R., Verkoczy, L., Wiehe, K., Munir Alam, S., Nicely, N.I., Santra, S., Bradley, T., Pemble, C.W., Zhang, J., Gao, F., et al. (2016). Initiation of immune tolerance–controlled HIV gp41 neutralizing B cell lineages. Sci. Transl. Med. 8, 336ra62–336ra62. 10.1126/scitranslmed.aaf0618.

62. Xu, K., Acharya, P., Kong, R., Cheng, C., Chuang, G.Y., Liu, K., Louder, M.K., O’Dell, S., Rawi, R., Sastry, M., et al. (2018). Epitope-based vaccine design yields fusion peptide-directed antibodies that neutralize diverse strains of HIV-1. Nat. Med. 24, 857–867. 10.1038/s41591-018-0042-6.

63. Kong, R., Duan, H., Sheng, Z., Xu, K., Acharya, P., Chen, X., Cheng, C., Dingens, A.S., Gorman, J., Sastry, M., et al. (2019). Antibody Lineages with Vaccine-Induced Antigen-Binding Hotspots Develop Broad HIV Neutralization. Cell 178, 567–584.e19. 10.1016/j.cell.2019.06.030.

64. Yang, Z., Dam, K.-M.A., Bridges, M.D., Hoffmann, M.A.G., DeLaitsch, A.T., Gristick, H.B., Escolano, A., Gautam, R., Martin, M.A., Nussenzweig, M.C., et al. (2022). Neutralizing antibodies induced in immunized macaques recognize the CD4-binding site on an occluded-open HIV-1 envelope trimer. Nat. Commun. 13, 732. 10.1038/s41467-022-28424-3.

65. Lynch, R.M., Wong, P., Tran, L., O’Dell, S., Nason, M.C., Li, Y., Wu, X., and Mascola, J.R. (2015). HIV-1 Fitness Cost Associated with Escape from the VRC01 Class of CD4 Binding Site Neutralizing Antibodies. J. Virol. 89, 4201–4213. 10.1128/jvi.03608-14.

66. Gorman, J., Soto, C., Yang, M.M., Davenport, T.M., Guttman, M., Bailer, R.T., Chambers, M., Chuang, G.-Y., DeKosky, B.J., Doria-Rose, N.A., et al. (2016). Structures of HIV-1 Env V1V2 with broadly neutralizing antibodies reveal commonalities that enable vaccine design. Nat. Struct. Mol. Biol. 23, 81–90. 10.1038/nsmb.3144.

67. Voss, J.E., Andrabi, R., Laura, E., Crispin, M., Wilson, I.A., Burton, D.R., Voss, J.E., Andrabi, R., Mccoy, L.E., Val, N.D., et al. (2017). Elicitation of Neutralizing Antibodies Targeting the V2 Apex of the HIV Envelope Trimer in a Wild-Type Animal Model. Cell Rep., 222–235.

68. Bonsignori, M., Hwang, K.-K., Chen, X., Tsao, C.-Y., Morris, L., Gray, E., Marshall, D.J., Crump, J.A., Kapiga, S.H., Sam, N.E., et al. (2011). Analysis of a Clonal Lineage of HIV-1 Envelope V2/V3 Conformational Epitope-Specific Broadly Neutralizing Antibodies and Their Inferred Unmutated Common Ancestors. J. Virol. 85, 9998–10009. 10.1128/jvi.05045-11.

69. Escolano, A., Gristick, H.B., Abernathy, M.E., Merkenschlager, J., Gautam, R., Oliveira, T.Y., Pai, J., West, A.P., Barnes, C.O., Cohen, A.A., et al. (2019). Immunization expands B cells specific to HIV-1 V3 glycan in mice and macaques. Nature 570, 468–473. 10.1038/s41586-019-1250-z.

70. Crooks, E.T., Osawa, K., Tong, T., Grimley, S.L., Dai, Y.D., Whalen, R.G., Kulp, D.W., Menis, S., Schief, W.R., and Binley, J.M. (2017). Effects of partially dismantling the CD4 binding site glycan fence of HIV-1 Envelope glycoprotein trimers on neutralizing antibody induction. Virology 505, 193–209. 10.1016/j.virol.2017.02.024.

71. Dosenovic, P., von Boehmer, L., Escolano, A., Jardine, J., Freund, N.T., Gitlin, A.D., McGuire, A.T., Kulp, D.W., Oliveira, T., Scharf, L., et al. (2015). Immunization for HIV-1 Broadly Neutralizing Antibodies in Human Ig Knockin Mice. Cell 161, 1505–1515. 10.1016/j.cell.2015.06.003.

72. Estes, J.D., Kityo, C., Ssali, F., Swainson, L., Makamdop, K.N., Del Prete, G.Q., Deeks, S.G., Luciw, P.A., Chipman, J.G., Beilman, G.J., et al. (2017). Defining total-body AIDS-virus burden with implications for curative strategies. Nat. Med. 23, 1271–1276. 10.1038/nm.4411.

73. Deleage, C., Immonen, T.T., Fennessey, C.M., Reynaldi, A., Reid, C., Newman, L., Lipkey, L., Schlub, T.E., Camus, C., O’Brien, S., et al. Defining early SIV replication and dissemination dynamics following vaginal transmission. Sci. Adv. 5, eaav7116. 10.1126/sciadv.aav7116.

74. Burton, D.R., and Mascola, J.R. (2015). Antibody responses to envelope glycoproteins in HIV-1 infection. Nat. Immunol. 16, 571–576. 10.1038/ni.3158.

75. Wang, H., Cheng, C., Dal Santo, J.L., Shen, C.-H., Bylund, T., Henry, A.R., Howe, C.A., Hwang, J., Morano, N.C., Morris, D.J., et al. Potent and broad HIV-1 neutralization in fusion peptide-primed SHIV-infected macaques. Cell. 10.1016/j.cell.2024.10.003.

76. Lynch Rebecca M., Tran Lillian, Louder Mark K., Schmidt Stephen D., Cohen Myron, DerSimonian Rebecca, Euler Zelda, Gray Elin S., Abdool Karim Salim, Kirchherr Jennifer, et al. (2012). The Development of CD4 Binding Site Antibodies during HIV-1 Infection. J. Virol. 86, 7588–7595. 10.1128/jvi.00734-12.

77. Gulla, K., Cibelli, N., Cooper, J.W., Fuller, H.C., Schneiderman, Z., Witter, S., Zhang, Y., Changela, A., Geng, H., Hatcher, C., et al. (2021). A non-affinity purification process for GMP production of prefusion-closed HIV-1 envelope trimers from clades A and C for clinical evaluation. Vaccine 39, 3379–3387. 10.1016/j.vaccine.2021.04.063.

78. Emsley, P., and Cowtan, K. (2004). ıt Coot: model-building tools for molecular graphics. Acta Crystallogr. Sect. D 60, 2126–2132. 10.1107/S0907444904019158.

79. Punjani, A., Rubinstein, J.L., Fleet, D.J., and Brubaker, M.A. (2017). cryoSPARC: algorithms for rapid unsupervised cryo-EM structure determination. Nat. Methods 14, 290–296. 10.1038/nmeth.4169.

80. Barad, B.A., Echols, N., Wang, R.Y.-R., Cheng, Y., DiMaio, F., Adams, P.D., and Fraser, J.S. (2015). EMRinger: side chain–directed model and map validation for 3D cryo-electron microscopy. Nat. Methods 12, 943–946. 10.1038/nmeth.3541.

81. Suloway, C., Pulokas, J., Fellmann, D., Cheng, A., Guerra, F., Quispe, J., Stagg, S., Potter, C.S., and Carragher, B. (2005). Automated molecular microscopy: The new Leginon system. J. Struct. Biol. 151, 41–60. 10.1016/j.jsb.2005.03.010.

82. Chen, V.B., Arendall, W.B., III, Headd, J.J., Keedy, D.A., Immormino, R.M., Kapral, G.J., Murray, L.W., Richardson, J.S., and Richardson, D.C. (2010). MolProbity: all-atom structure validation for macromolecular crystallography. Acta Crystallogr. Sect. D 66, 12–21. 10.1107/S0907444909042073.

83. Adams, P.D., Gopal, K., Grosse-Kunstleve, R.W., Hung, L.-W., Ioerger, T.R., McCoy, A.J., Moriarty, N.W., Pai, R.K., Read, R.J., Romo, T.D., et al. (2004). Recent developments in the ıt PHENIX software for automated crystallographic structure determination. J. Synchrotron Radiat. 11, 53–55. 10.1107/S0909049503024130.

84. Krissinel, E., and Henrick, K. (2007). Inference of Macromolecular Assemblies from Crystalline State. J. Mol. Biol. 372, 774–797. 10.1016/j.jmb.2007.05.022.

85. Letvin, N.L., and King, N.W. (1990). Immunologic and Pathologic Manifestations of the Infection of Rhesus Monkeys with Simian Immunodeficiency Virus of Macaques. JAIDS J. Acquir. Immune Defic. Syndr. 3.

86. Landais, E., Huang, X., Havenar-Daughton, C., Murrell, B., Price, M.A., Wickramasinghe, L., Ramos, A., Bian, C.B., Simek, M., Allen, S., et al. (2016). Broadly Neutralizing Antibody Responses in a Large Longitudinal Sub-Saharan HIV Primary Infection Cohort. PLoS Pathog. 12, 1–22. 10.1371/journal.ppat.1005369.

87. Sarzotti-Kelsoe, M., Bailer, R.T., Turk, E., Lin, C., Bilska, M., Greene, K.M., Gao, H., Todd, C.A., Ozaki, D.A., Seaman, M.S., et al. (2014). Optimization and validation of the TZM-bl assay for standardized assessments of neutralizing antibodies against HIV-1. J. Immunol. Methods 409, 131–146. 10.1016/j.jim.2013.11.022.

88. Seaman, M.S., Janes, H., Hawkins, N., Grandpre, L.E., Devoy, C., Giri, A., Coffey, R.T., Harris, L., Wood, B., Daniels, M.G., et al. (2010). Tiered Categorization of a Diverse Panel of HIV-1 Env Pseudoviruses for Assessment of Neutralizing Antibodies. J. Virol. 84, 1439–1452. 10.1128/jvi.02108-09.

89. deCamp, A., Hraber, P., Bailer, R.T., Seaman, M.S., Ochsenbauer, C., Kappes, J., Gottardo, R., Edlefsen, P., Self, S., Tang, H., et al. (2014). Global Panel of HIV-1 Env Reference Strains for Standardized Assessments of Vaccine-Elicited Neutralizing Antibodies. J. Virol. 88, 2489–2507. 10.1128/jvi.02853-13.

90. Perfetto, S.P., Chattopadhyay, P.K., Lamoreaux, L., Nguyen, R., Ambrozak, D., Koup, R.A., and Roederer, M. (2010). Amine-Reactive Dyes for Dead Cell Discrimination in Fixed Samples. Curr. Protoc. Cytom. 53, 9.34.1–9.34.14. 10.1002/0471142956.cy0934s53.

91. Donaldson, M.M., Kao, S.-F., Eslamizar, L., Gee, C., Koopman, G., Lifton, M., Schmitz, J.E., Sylwester, A.W., Wilson, A., Hawkins, N., et al. (2012). Optimization and qualification of an 8-color intracellular cytokine staining assay for quantifying T cell responses in rhesus macaques for pre-clinical vaccine studies. J. Immunol. Methods 386, 10–21. 10.1016/j.jim.2012.08.011.

92. Huang, J., Doria-Rose, N.A., Longo, N.S., Laub, L., Lin, C.-L., Turk, E., Kang, B.H., Migueles, S.A., Bailer, R.T., Mascola, J.R., et al. (2013). Isolation of human monoclonal antibodies from peripheral blood B cells. Nat. Protoc. 8, 1907–1915. 10.1038/nprot.2013.117.

93. Doria-Rose, N., Doria-Rose, N., Bailer, R., Louder, M., Lin, C.-L., Turk, E., Laub, L., Longo, N., Connors, M., and Mascola, J. (2013). High throughput HIV-1 microneutralization assay. Protoc. Exch. 10.1038/protex.2013.069.

94. Liao, H.-X., Levesque, M.C., Nagel, A., Dixon, A., Zhang, R., Walter, E., Parks, R., Whitesides, J., Marshall, D.J., Hwang, K.-K., et al. (2009). High-throughput isolation of immunoglobulin genes from single human B cells and expression as monoclonal antibodies. J. Virol. Methods 158, 171–179. 10.1016/j.jviromet.2009.02.014.

95. Smith, K., Garman, L., Wrammert, J., Zheng, N.-Y., Capra, J.D., Ahmed, R., and Wilson, P.C. (2009). Rapid generation of fully human monoclonal antibodies specific to a vaccinating antigen. Nat. Protoc. 4, 372–384. 10.1038/nprot.2009.3.

96. Tiller, T., Meffre, E., Yurasov, S., Tsuiji, M., Nussenzweig, M.C., and Wardemann, H. (2008). Efficient generation of monoclonal antibodies from single human B cells by single cell RT-PCR and expression vector cloning. J. Immunol. Methods 329, 112–124. 10.1016/j.jim.2007.09.017.

97. Sanders, R.W., Derking, R., Cupo, A., Julien, J.-P., Yasmeen, A., de Val, N., Kim, H.J., Blattner, C., de la Peña, A.T., Korzun, J., et al. (2013). A Next-Generation Cleaved, Soluble HIV-1 Env Trimer, BG505 SOSIP.664 gp140, Expresses Multiple Epitopes for Broadly Neutralizing but Not Non-Neutralizing Antibodies. PLOS Pathog. 9, 1–20. 10.1371/journal.ppat.1003618.

98. Pancera, M., Zhou, T., Druz, A., Georgiev, I.S., Soto, C., Gorman, J., Huang, J., Acharya, P., Chuang, G.-Y., Ofek, G., et al. (2014). Structure and immune recognition of trimeric pre-fusion HIV-1 Env. Nature 514, 455–461. 10.1038/nature13808.

99. Kwon, Y.D., Chuang, G.-Y., Zhang, B., Bailer, R.T., Doria-Rose, N.A., Gindin, T.S., Lin, B., Louder, M.K., McKee, K., O’Dell, S., et al. (2018). Surface-Matrix Screening Identifies Semi-specific Interactions that Improve Potency of a Near Pan-reactive HIV-1-Neutralizing Antibody. Cell Rep. 22, 1798–1809. 10.1016/j.celrep.2018.01.023.

100. Voss, N.R., Yoshioka, C.K., Radermacher, M., Potter, C.S., and Carragher, B. (2009). DoG Picker and TiltPicker: Software tools to facilitate particle selection in single particle electron microscopy. J. Struct. Biol. 166, 205–213. 10.1016/j.jsb.2009.01.004.

101. Lander, G.C., Stagg, S.M., Voss, N.R., Cheng, A., Fellmann, D., Pulokas, J., Yoshioka, C., Irving, C., Mulder, A., Lau, P.-W., et al. (2009). Appion: An integrated, database-driven pipeline to facilitate EM image processing. J. Struct. Biol. 166, 95–102. 10.1016/j.jsb.2009.01.002.

102. Zheng, S.Q., Palovcak, E., Armache, J.-P., Verba, K.A., Cheng, Y., and Agard, D.A. (2017). MotionCor2: anisotropic correction of beam-induced motion for improved cryo-electron microscopy. Nat. Methods 14, 331–332. 10.1038/nmeth.4193.

103. Rohou, A., and Grigorieff, N. (2015). CTFFIND4: Fast and accurate defocus estimation from electron micrographs. J. Struct. Biol. 192, 216–221. 10.1016/j.jsb.2015.08.008.

104. Zhang, K. (2016). Gctf: Real-time CTF determination and correction. J. Struct. Biol. 193, 1–12. 10.1016/j.jsb.2015.11.003.

105. Scheres, S.H.W. (2012). RELION: Implementation of a Bayesian approach to cryo-EM structure determination. J. Struct. Biol. 180, 519–530. 10.1016/j.jsb.2012.09.006.

106. Davis, I.W., Murray, L.W., Richardson, J.S., and Richardson, D.C. (2004). MolProbity: structure validation and all-atom contact analysis for nucleic acids and their complexes. Nucleic Acids Res. 32, W615–W619. 10.1093/nar/gkh398.

107. Krebs, S.J., Kwon, Y.D., Schramm, C.A., Law, W.H., Donofrio, G., Zhou, K.H., Gift, S., Dussupt, V., Georgiev, I.S., Schätzle, S., et al. (2019). Longitudinal Analysis Reveals Early Development of Three MPER-Directed Neutralizing Antibody Lineages from an HIV-1-Infected Individual. Immunity 50, 677–691.e13. 10.1016/j.immuni.2019.02.008.

108. Ramesh, A., Darko, S., Hua, A., Overman, G., Ransier, A., Francica, J.R., Trama, A., Tomaras, G.D., Haynes, B.F., Douek, D.C., et al. (2017). Structure and diversity of the rhesus macaque immunoglobulin loci through multiple de novo genome assemblies. Front. Immunol. 8. 10.3389/fimmu.2017.01407.

109. Hraber, P., Korber, B., Wagh, K., Giorgi, E.E., Bhattacharya, T., Gnanakaran, S., Lapedes, A.S., Learn, G.H., Kreider, E.F., Li, Y., et al. (2015). Longitudinal antigenic sequences and sites from intra-host evolution (LASSIE) identifies immune-selected HIV variants. Viruses 7, 5443–5475. 10.3390/v7102881.

110. Hoffenberg, S., Powell, R., Carpov, A., Wagner, D., Wilson, A., Pond, S.K., Lindsay, R., Arendt, H., DeStefano, J., Phogat, S., et al. (2013). Identification of an HIV-1 Clade A Envelope That Exhibits Broad Antigenicity and Neutralization Sensitivity and Elicits Antibodies Targeting Three Distinct Epitopes. J. Virol. 87, 5372–5383. 10.1128/jvi.02827-12.

